# *In situ* deposition of nanobodies by an engineered commensal microbe promotes survival in a mouse model of enterohemorrhagic *E. coli*

**DOI:** 10.1101/2024.07.30.605899

**Authors:** Rajkamal Srivastava, Coral González-Prieto, Jason P Lynch, Michele Muscolo, Catherine Y Lin, Markus A Brown, Luisa Lemos, Anishma Shrestha, Marcia S Osburne, John M Leong, Cammie F Lesser

## Abstract

Engineered smart microbes that deliver therapeutic payloads are emerging as treatment modalities, particularly for diseases with links to the gastrointestinal tract. Enterohemorrhagic *E coli* (EHEC) is a causative agent of potentially lethal hemolytic uremic syndrome. Given concerns that antibiotic treatment increases EHEC production of Shiga toxin (Stx), which is responsible for systemic disease, novel remedies are needed. EHEC encodes a type III secretion system (T3SS) that injects Tir into enterocytes. Tir inserts into the host cell membrane, exposing an extracellular domain that subsequently binds intimin, one of its outer membrane proteins, triggering the formation of attaching and effacing (A/E) lesions that promote EHEC mucosal colonization. *Citrobacter rodentium* (Cr), a natural A/E mouse pathogen, similarly requires Tir and intimin for its pathogenesis. Mice infected with Cr(ΦStx2dact), a variant lysogenized with an EHEC-derived phage that produces Stx2dact, develop intestinal A/E lesions and toxin-dependent disease. Stx2a is more closely associated with human disease. By developing an efficient approach to seamlessly modify the *C. rodentium* genome, we generated Cr_Tir-M^EHEC^(ΦStx2a), a variant that expresses Stx2a and the EHEC extracellular Tir domain. We found that mouse pre-colonization with HS-PROT_3_EcT-TD4, a human commensal *E. coli* strain (*E. coli* HS) engineered to efficiently secrete- an anti-EHEC Tir nanobody, delayed bacterial colonization and improved survival after challenge with Cr_Tir-M^EHEC^(ΦStx2a). This study provides the first evidence to support the efficacy of engineered commensal *E. coli* to intestinally deliver therapeutic payloads that block essential enteric pathogen virulence determinants, a strategy that may serve as an antibiotic-independent antibacterial therapeutic modality.

**Significance Statement:** Engineered smart microbes that secrete therapeutics are emerging as treatment modalities, particularly for gut-based diseases. With the growing threat of multidrug-resistant infection, non-antibiotic treatments are urgently needed. The gastrointestinal pathogen enterohemorrhagic *E coli* (EHEC) can cause the potentially lethal hemolytic uremic syndrome, a toxin-driven disease. Given concerns that antibiotics increase toxin release, treatment is largely limited to supportive care. Here, we show that pre-treatment with a commensal *E. coli* (HS-PROT_3_EcT) engineered to secrete an antibody that blocks an essential EHEC virulence factor delays the establishment of an EHEC-like infection in mice. This study strongly suggests that smart microbes that deliver payloads that block colonization factors of gut pathogens can be developed as critically needed alternatives to antibiotics for fighting bacterial infections.

## Introduction

Smart microbes that deliver therapeutic payloads are emerging as new treatment modalities for a variety of pathologies, particularly those with links to the gastrointestinal tract^1–3^. By acting specifically at sites of diseases, these microbes provide a means to improve therapeutic efficacy while decreasing off-target side effects. Various chassis are being explored, including yeast, Gram-positive and Gram-negative bacteria.

In the case of Gram-positive bacteria, their native secretion systems have been repurposed to deliver therapeutic protein payloads into their surroundings. However, similar approaches are limited for the more genetically tractable Gram-negative bacteria, which, in contrast to the Gram-positive bacteria, at least transiently colonize the intestines^4,5^. This is because their native secretion systems primarily target the delivery of proteins into the periplasm, the region between their inner and outer membranes, or onto their outer envelope. To circumvent this limitation of Gram-negative bacteria, investigators have designed variants of *E. coli* that release therapeutic payloads by lysis^6,7^ or display proteins of interest on their surface via their fusion to outer surface proteins^8,9^.

T3SSs are complex nanomachines common to Gram-negative bacterial pathogens that function to directly transport tens of proteins, commonly referred to as effectors, directly into the cytosol of targeted host cells. These complex machines, composed of >20 different proteins, are embedded within their outer bacterial envelope with a needle-like extension^10,11^. The needle is capped with a tip complex that holds the machine in an off configuration^12^. Upon contact with host cells, the tip complex forms pores in the host cell membrane, triggering and enabling the direct injection of effectors into the host cell cytosol. In the absence of the tip complex, the modified nanomachine constitutively secretes proteins into its surroundings.

Leveraging our expertise in bacterial secretion systems, we previously developed the PROT_3_EcT (<u>PRO</u>biotic <u>T</u>ype <u>3</u> secretion *<u>E</u>. <u>c</u>oli* <u>T</u>herapeutic) platform, a suite of engineered probiotic and commensal *E. coli* that express a modified bacterial type III secretion system (T3SS) that lacks the tip complex and secretes defined therapeutic payloads into its surroundings^13,14^. This platform is modular in design such that the *E. coli* chassis, protein payloads, and their modes of transcriptional regulation can be easily exchanged or modified^13,14^.

Enterohemorrhagic *E. coli* (EHEC) is a food-borne pathogen that causes bloody diarrhea and, in some cases, life-threatening hemolytic uremic syndrome (HUS), a clinical syndrome defined by the triad of hemolytic anemia, thrombotic thrombocytopenia, and renal failure. EHEC, together with enteropathogenic *E coli* (EPEC) and *Citrobacter rodentium* (Cr), form the family of attaching/effacing (A/E) pathogens. EPEC and EHEC are human-specific pathogens, while Cr targets mice. The attachment of each of these pathogens to intestinal epithelial cells is dependent on a highly conserved T3SS^15^. The first effector delivered by this T3SS into host cells is Tir (translocated intimin receptor).

Upon entry, Tir inserts into the mammalian plasma membrane, exposing on the surface of the host cells a domain, Tir-M, that serves as the receptor for intimin^16,17^, a bacterial outer membrane protein common to A/E pathogens. Intimin binding leads to a higher-order clustering of Tir and the assembly of actin pedestals beneath the attached bacteria, an essential step in the pathogenesis of A/E pathogens^18,19^.

EHEC is unique in its ability to induce HUS due to its secretion of Shiga toxins (Stx). Two main types of Stx exist: Stx1 and Stx2. Stx2 is associated with more severe disease^20^. There are at least seven variants of Stx2. Stx2dact is uniquely proteolytically activated by elastase in the intestinal mucosa and has been associated with mouse virulence^21^, whereas Stx2a is most often associated with human disease^20^. No vaccines or specific therapeutic interventions are currently available to prevent or treat EHEC. Antibiotic treatments can lead to a bacterial SOS response that triggers phage induction and increased Stx release^22^.

Towards developing a new approach for the treatment of EHEC, Ruano-Gallego and colleagues isolated a neutralizing camelid-derived single domain ant-Tir antibody (nanobody) (Nb^TD4^). They found that Nb^TD4^ binds with high affinity specifically to EHEC Tir, blocking its interaction with intimin^23^. They discovered that Nb^TD4^ blocks the formation of EHEC actin pedestals on cell lines or human colonic biopsies, but did not investigate whether Nb^TD4^ blocks the establishment of an *in vivo* infection.

Efficient colonization of mice by EHEC requires the infection of germ-free mice or mice pretreated with antibiotics. These mice succumb to Stx-dependent disease^24^, but factors needed for the formation of actin pedestals, such as Tir and intimin, play no role in these infection models^25,26^. In contrast, conventional mice are efficiently colonized by Cr, which generates A/E lesions in a Tir/intimin-dependent manner that are essential for intestinal colonization and the development of disease^27^. As Cr does not encode Stx, to model A/E lesion formation and Stx-mediated systemic disease, Mallick and colleagues lysogenized Cr with a Stx-producing phage (ΦStx2dact) derived from a naturally occurring EHEC strain^28^. Mice inoculated with the resulting strain Cr(ΦStx2dact) develop lethal disease featuring weight loss, intestinal inflammation, renal pathology, and proteinuria. However, several key virulence factors of Cr(ΦStx2dact) vary from their EHEC counterpart, and in human clinical isolates of EHEC, Stx2dact is less common than Stx2a.

Our goal in this study was to test the potential efficacy of Nb^TD4^-secreting PROT_3_EcT in a model that closely resembles EHEC infection and disease. We developed an efficient approach to seamlessly generate Cr_Tir-M^EHEC^(ΦStx2a), a *C. rodentium* strain that produces Stx2a rather than Stx2dact, and a chimeric Cr/EHEC Tir with the extracellular EHEC Tir-M domain, thus a more humanized strain. This completely recombination-based approach can likely be extended to all genetically tractable enteric Gram-negative bacteria, including pathogens and laboratory strains of *E. coli*.

In parallel, we developed HS-PROT_3_EcT-TD4, an *E. coli* HS variant of PROT_3_EcT engineered to secrete the anti-Tir Nb^TD4^ via a modified type III secretion sequence. Thus, demonstrating the modularity of the PROT_3_EcT platform, in terms of bacterial chassis and therapeutic payloads. Strikingly, we found that mice pre-colonized with HS-PROT_3_EcT-TD4 displayed delayed Cr_Tir-M^EHEC^(ΦStx2a) colonization and survived for 3-4 days longer than mice that were mock-treated or colonized with an HS-PROT_3_EcT strain that does not secrete the nanobody. This study provides the first evidence to support the efficacy of engineered commensal *E. coli* to intestinally deliver therapeutic payloads that block essential enteric pathogen virulence determinants, a strategy that may serve as an antibiotic-independent antibacterial therapeutic modality.

## Results and Discussion

### Development of an efficient seamless cloning approach for Cr

Cr has been utilized to model human infection by EPEC (enteropathogenic *E. coli*) due to its ability to generate A/E lesions on intestinal epithelium. Upon infection with native Cr, mice develop modest and transient weight loss and diarrhea accompanied by histological evidence of epithelial crypt hyperplasia in the colon. Bacterial colonization is maximal at 7-10 days post-inoculation and is usually cleared within 3-4 weeks^29^. Cr does not produce Shiga toxin, so Cr(ΦStx2dact) was developed to model both colonization and toxigenic aspects of human EHEC infections^30^. Mice infected with this strain succumb to disease within 7-10 days and, on necropsy, demonstrate evidence of renal damage^28,31^.

To further expand its applicability to EHEC disease, we developed an efficient recombination-based platform to modify Cr(ΦStx2dact) by seamlessly swapping regions of Cr chromosomal DNA with those of its EHEC homologs. This experimental pipeline incorporates aspects of site-specific^32^ and homologous recombination^33^ plus the introduction of precise gaps in chromosomal DNA via the introduction of I-SceI recognition sites, an 18-base pair recognition site not naturally found in the genome of Cr and other related species^34^ (Fig. 1A).

**Figure 1:**
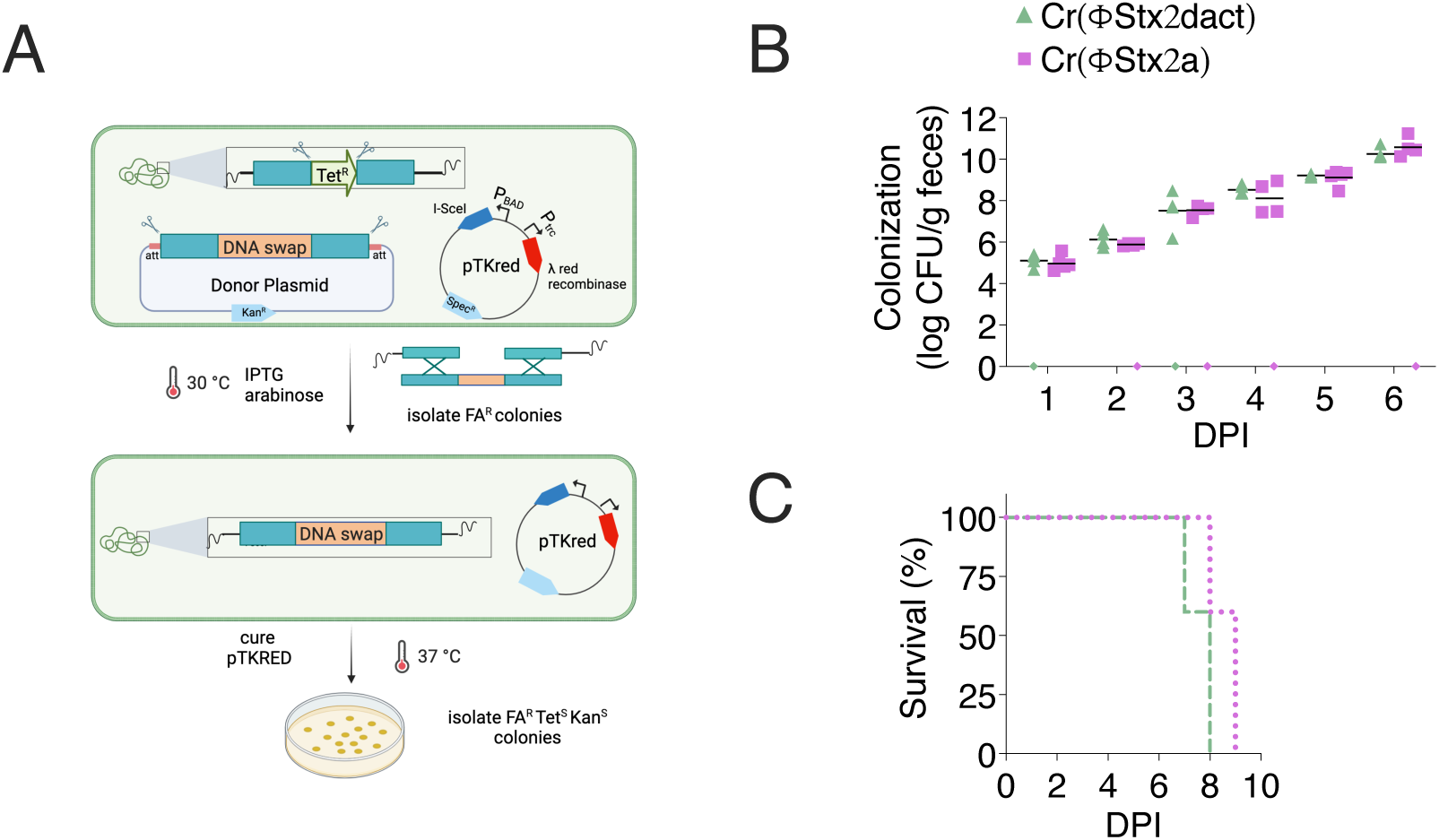
Mice are equally susceptible to infection with Cr(ΦStx2dact) or Cr(ΦStx2a). (A) Schematic of seamless cloning approach. (B, C) Seven-week-old female mice were orally inoculated with 1 × 10^8^ CFU of Cr(ΦStx2dact) or Cr(ΦStx2a) by feeding. Five mice were included in each cohort. (B) Viable counts of bacteria in feces were determined by plating. Each point shown represents an individual mouse, and each line represents the geometric mean of log^10^ CFU/g of feces. Samples plotted on the x-axis indicate no data available due to a lack of collected feces. All differences in bacterial titers were ns (non-significant) as determined by two-way ANOVA. (C) Kaplan–Meier survival curves of mice infected with Cr(ΦStx2dact) and Cr(ΦStx2a). No evidence of statistical significance between the two cohorts was found using the log-rank (Mantel-Cox) test.

In the first phase of this pipeline, homologous recombination is used to replace the region in the Cr genome to be swapped with a tetracycline resistance (Tet^R^) cassette flanked on each side by I-SceI sites. In parallel, the DNA sequence to be introduced, flanked on each side by ∼300 base pairs of homology and outer I-SceI and attB recognition sites, is introduced onto a kanamycin-resistant Gateway entry vector. This resulting plasmid is referred to as the “donor plasmid.” Once both these steps are completed, the Tet^R^-containing strain is transformed with the donor plasmid and pTKRED, a spectinomycin-resistant temperature-sensitive (ts) plasmid that encodes IPTG-inducible *λ*-red recombinase and arabinose-inducible I-SceI^34^.

In the second phase of the pipeline, the transformed bacteria are grown in the presence of arabinose and IPTG (isopropyl ß-D-1-thiogalactopyranoside) to induce expression of I-SceI and the λ-red recombinase, respectively. One then screens for colonies sensitive to tetracycline and kanamycin, which have recombined the sequence of interest onto the chromosome, followed by curing on pTKRED and loss of spectinomycin resistance.

When optimizing the protocol, we found that Cr arabinose induction is more robust in minimal (M9/0.5% glycerol) as compared to rich (LB or SOB) media (Fig. S1) and that tetracycline-sensitive (Tet^S^) colonies are enriched when using fusaric-acid as a counter-selection (Fig. S2).

### Mice exhibit similar patterns of susceptibility to Stx2dact and Stx2a

Shiga toxin, the key virulence factor in systemic EHEC disease, consists of two subunits. The A subunit of Stx2 is an N-glycosidase that cleaves and inactivates the 28s RNA subunit of the 60S ribosome, thus blocking translation. The B subunit forms a pentamer that binds to Gb3, the cellular toxin receptor that is primarily found on endothelial cells. Both subunits are encoded in an operon located within an inducible lysogenic λ-like bacteriophage^20^. Although the A and B subunits of Stx2a and Stx2dact share a high degree of identity, differences are found within both subunits (Fig. S3), and Stx2a is the most prevalent variant in patients who develop HUS^35^. As a first test of the seamless cloning pipeline, we swapped the region of DNA containing the genes encoding both subunits of Stx2dact with the equivalent sequences encoding Stx2a to generate Cr(ΦStx2a).

After confirming that the *in vitro* growth rates of Cr(ΦStx2dact) and Cr(ΦStx2a) were indistinguishable (Fig. S4A), we compared the fate of mice infected with each strain. We used the food inoculation model, which was previously established to lead to a highly synchronized infection^31^. C57BL/6 mice starved overnight were fed a small fragment of chow carrying ∼10^8^ colony forming units (CFU) of each strain. We observed similar intestinal expansion kinetics of Cr(ΦStx2dact) and Cr(ΦStx2a), as assessed by daily quantitation of the colony counts found in shed feces (Fig. 1B). Mice infected with each strain exhibited very similar survival curves (Fig. 1C). Thus, at least when administered at 10^8^ CFU, mice are equally susceptible to infection with Cr(ΦStx2dact) and Cr(ΦStx2a).

### The extracellular Tir-M domains of Cr and EHEC are functionally interchangeable

The pathogenesis of A/E pathogens is dependent on their highly conserved T3SSs^36^. The first effector injected by each into host cells is Tir^37^. After entering the host cell cytosol, Tir inserts into the plasma membrane in a hairpin-loop conformation with an extracellular Tir-M domain flanked by cytosolic N- and C-terminal domains (Fig. 2A). Tir-M binds to intimin, leading to the assembly of actin pedestals beneath the attached bacteria.

**Figure 2:**
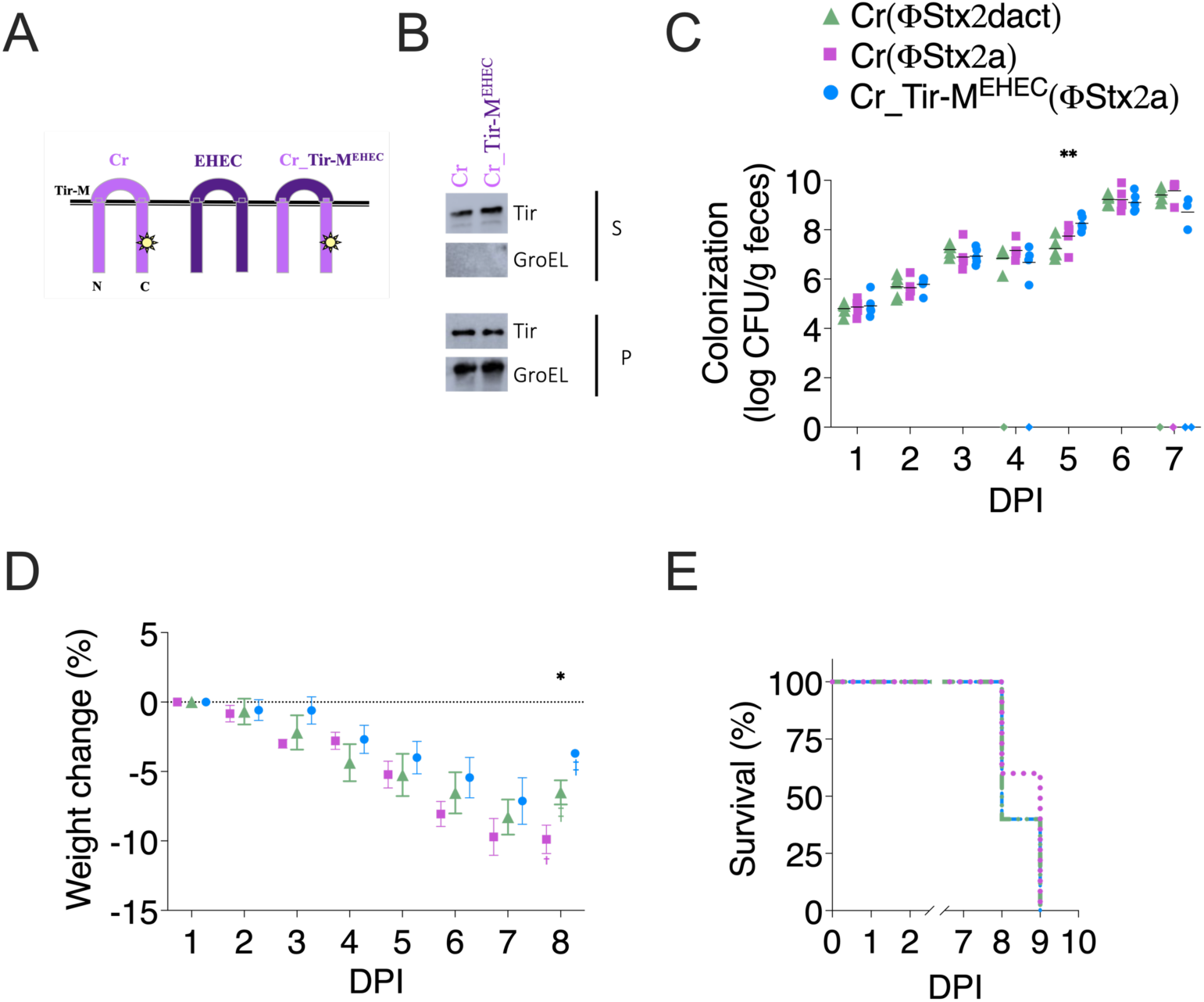
The extracellular domains of EHEC, EPEC, and Cr Tir are functionally interchangeable. (A) Schematic of WT and chimeric Tir variants. (B) Secretion assay of Tir variants from designated strains. Supernatants (S) and whole-cell pellet lysates (P) fractions were obtained 6 hours post-transfer to fresh media. Images of immunoblots probed with an anti-Tir or an anti-GroEL antibody are shown. GroEL serves as a lysis control for (S) and loading control for (P). (C-E). Seven-week-old female mice were orally inoculated with 1 × 10^8^ CFU designated Cr strains. Five mice were included in each cohort. (C) Viable counts of bacteria in feces were determined by plating. Each point shown represents an individual mouse, and each line represents the geometric mean of log^10^ CFU/g of feces. Samples plotted on the x-axis indicate no data available due to a lack of collected feces. (D) Time course of body weight changes (%) over time. Mean +/-SEM plotted. Data in C and D were analyzed using two-way ANOVA with Bonferroni’s post hoc multiple comparison test at a 95% confidence interval. For C, DPI (1-4, 6, 7) = ns; DPI 5, Cr(ΦStx2dact) vs Cr_Tir-M^EHEC^(ΦStx2a) **p =0.0036. For D, DPI (1-7) = ns, DPI 8, Cr(ΦStx2a) vs Cr_Tir-M^EHEC^(ΦStx2a) *p = 0.0135. (E) Kaplan–Meier survival curves of mice infected with Cr(ΦStx2dact), Cr(ΦStx2a), and Cr_Tir-M^EHEC^(ΦStx2a). No evidence of statistical significance was found between the four cohorts using the log-rank (Mantel-Cox) test.

The pedestal generating mechanisms of Cr and EHEC Tir differ^38^. Upon binding intimin, a tyrosine in the C-terminal Cr Tir domain is phosphorylated, an event sufficient to recruit the actin assembly machinery^27,39^. EHEC Tir lacks this tyrosine^26,40^. The formation of EHEC pedestals is dependent on EspFu (also known as TccP)^41,42^, an EHEC-specific type III secreted effector. EspFu is required for the formation of the EHEC Tir-containing actin assembly complex needed for the generation of pedestals. These differences in the C-termini of Cr and EHEC Tir likely explain why Cr and EHEC Tir are not functionally interchangeable^43,44^.

The Tir-M domains of EHEC and Cr share a high degree of homology, and Cr intimin can bind to EHEC Tir^45^. Yet, Ruano-Gallego and colleagues found that Nb^TD4^, the Nb that blocks Tir binding to intimin, binds much more strongly to the Tir-M domain of EHEC than that of Cr, likely due to a few amino acid differences^23^. With the goal of investigating whether secreted Nb^TD4^ can block an *in vivo* infection, we first tested whether, by swapping the Tir-M domains of Cr and EHEC, we could generate Stx-producing Cr variants with EHEC characteristics that would still cause disease in mice.

Using the seamless cloning approach, we swapped the Tir-M domain of Cr with that of EHEC in both the CrΦStx2a and CrΦStx2dact backgrounds, resulting in strains referred to as Cr_Tir-M^EHEC^(ΦStx2a) and Cr_Tir-M^EHEC^(ΦStx2dact). Regardless of which Tir or Stx2 they encode, each strain exhibited similar *in vitro* growth rates (Fig. S4B-C).

Furthermore, we found that Cr(ΦStx2a) and Cr_Tir-M^EHEC^(ΦStx2a) secreted equivalent levels of Tir (Fig. 2B). This was expected, as the sequences that define Tir as a secreted protein are all contained in its first N-terminal 80 residues, a region upstream of Tir-M^46^.

We next compared the fate of C57BL/6 mice infected with ∼10^8^ CFU of Cr(ΦStx2dact), Cr(ΦStx2a), and the further “humanized” Cr_Tir-M^EHEC^(ΦStx2a) via the food-borne inoculation route. We observed no differences in the kinetics of colonization of the strains as assessed by fecal shedding (Fig. 2C), with titers for each strain increasing over time. Mice infected with each strain exhibited similar patterns of weight loss (Fig. 2D), and all succumbed to the infection on day 8 or 9 (Fig. 2E). These observations demonstrate that the extracellular Tir-M domains of Cr and EHEC are functionally interchangeable. Thus, we have expanded the variants of Cr(ΦStx2) that can be used to monitor aspects of infection specific to the EHEC Tir-M domain.

### EcN-PROT_3_EcT delays the susceptibility of mice to Cr(ΦStx2dact)

T3SSs like those present in *Shigella* and A/E pathogens function to inject effectors directly into targeted host cells. However, when the proteins that form the tip complex that holds the machine in an off configuration prior to host cell contact are removed, the machine constitutively secretes proteins into its surroundings^12^. We previously established that when this modified *Shigella* T3SS is introduced into non-pathogenic laboratory and probiotic *E. coli,* including Nissle 1917 *E. coli* (EcN). The strains robustly and constitutively secrete proteins, including functional nanobodies, into their surroudings^13^. EcN outfitted with this secretion system is referred to herein as EcN-PROT_3_EcT (PRObiotic Type III secretion *E. coli* Therapeutic).

Under its tradename of Mutaflor, EcN is used in Europe and Canada for the treatment of IBD due to its inherent anti-inflammatory activities^47^. EcN also has antibacterial activities^5,48,49^, including blocking EHEC colonization of mice^50^. Thus, before testing whether EcN-PROT_3_EcT secreted anti-Tir TD4 nanobodies would block infection with Cr, we investigated whether EcN-PROT_3_EcT would block infection with Cr(çStx2dact).

C56BL/6 mice were orally inoculated with three doses of 10^9^ CFU of EcN-PROT_3_EcT or diluent (20% sucrose) at 4-5 day intervals, reaching a stable level of colonization reflective of the shedding of ∼10^5^ CFU/g of feces (Fig. S5A). Subsequently, the mice were orally inoculated with 10^8^ CFU of Cr(ΦStx2dact). Mice pre-colonized with EcN-PROT_3_EcT demonstrated delayed colonization (Fig. 3A) and weight loss (Fig. 3B) and survived 4-5 days longer than those that previously solely received the diluent (Fig. 3C). The determinants that enable EcN to delay Cr colonization remain to be characterized. Nevertheless, given EcN’s inherent anti-Cr activity, we investigated the use of another *E. coli* strain as our PROT_3_EcT chassis.

**Figure 3:**
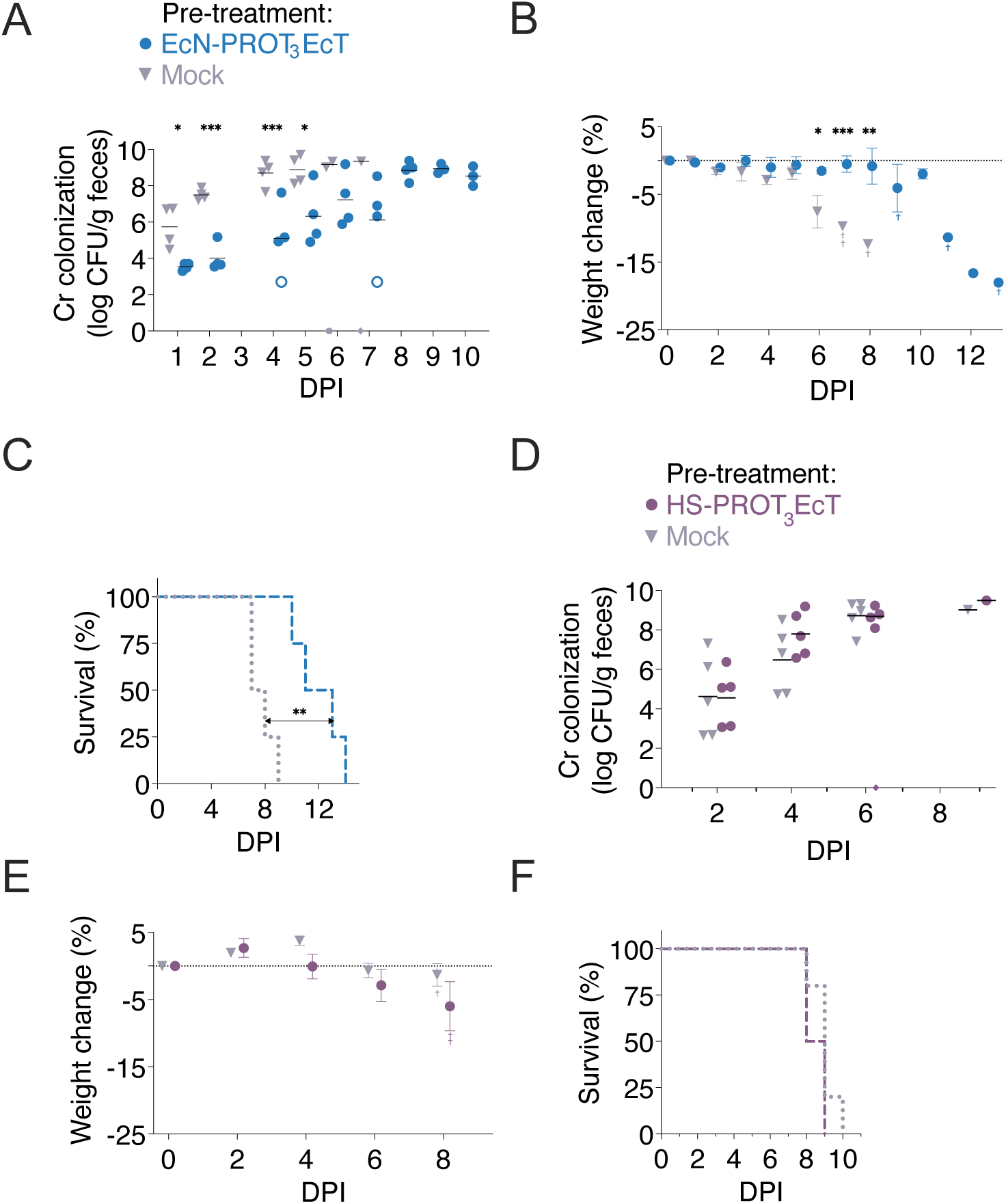
EcN-but not HS-PROT_3_EcT significantly delays infection with Cr(ΦStx2). Six-seven-week-old female mice pre-treated with (A-C) EcN-PROT_3_EcT or (D-F) HS-PROT_3_EcT or (A-F) mock media were infected with 1×10^8^ CFU of Cr(ΦStx2dact) (A-C) or Cr(ΦStx2a) (D-F). Five mice were included in each cohort. (A, D) Viable counts of bacteria in feces were determined by plating. Each point shown represents an individual mouse, and each line represents the geometric mean. Samples plotted on the x-axis indicate no data available. Open symbols indicate CFU at the limit of detection (LOD). This value was used when evaluating statistical significance at each time point using two-way ANOVA with Bonferroni’s post hoc multiple comparison test (95% CI). In (A), DPI 1: p = 0.0414; DPI 2: p = 0.0007, DPI 4: p = 0.0005; DPI 5: p = 0.0138; DPI 6-7: ns. In (D), all time points were found to be ns. (B, E) Time course of body weight changes (%) over time. Mean +/-SEM plotted. A two-tailed unpaired Student’s t-test was used to determine statistical significance (95% CI). In (B), DPI 1-5: ns; DPI 6: p = 0.0492, DPI 7: p = 0.0003, DPI 8: 0.0047. In (E), all differences were ns. (C, F) Kaplan–Meier survival curves of mice group pre-treated with designated strains infected with Cr(ΦStx2). Statistical significance was determined by the log-rank (Mantel-Cox) test. Differences in survival in (C) (P = 0.0062) but not (F) were found to be statistically significant.

### HS-PROT_3_EcT stably colonizes mice but does not block infection with Cr(ΦStx2a)

*E. coli* HS is a human commensal^51^ previously established to at least transiently colonize the intestines of mice^50,52^. Pre-colonization with *E. coli* HS was previously found not to block EHEC colonization^50^. In prior studies, we demonstrated that the modified *Shigella* T3SS is functional when introduced into *E. coli* HS, at least when the operons encoding the modified T3SS were inserted onto the chromosome, and their shared transcription regulator, VirB, expressed from an IPTG-inducible Ptrc promoter on a plasmid maintained via antibiotic resistance^13^.

For animal studies, to avoid a requirement for antibiotics for the maintenance of the plasmids encoding VirB as well as a therapeutic payload like Nb^TD4^, we performed the following modifications to the *E. coli* HS PROT_3_EcT variant. First, we introduced a gene cassette that encodes *virB,* controlled by a constitutive promoter (PJ23119), onto the chromosome. Second, we deleted the *E. coli* HS genes that encode its two functionally redundant alanine racemases, *alr* and *dadX,* to generate a variant that would maintain a plasmid via auxotrophic selection. Alanine racemases act to convert L-ala to D-ala, an essential cell wall component that is present at insufficient levels in the mammalian intestines to support *E. coli* growth^53^. This strain, referred to as HS-PROT_3_EcT, can be maintained on media supplemented with D-alanine or when transformed with an *alr*-encoding plasmid. All references to HS-PROT_3_EcT herein refer to a variant transformed with an *alr*-containing plasmid (Fig. 4A).

**Figure 4:**
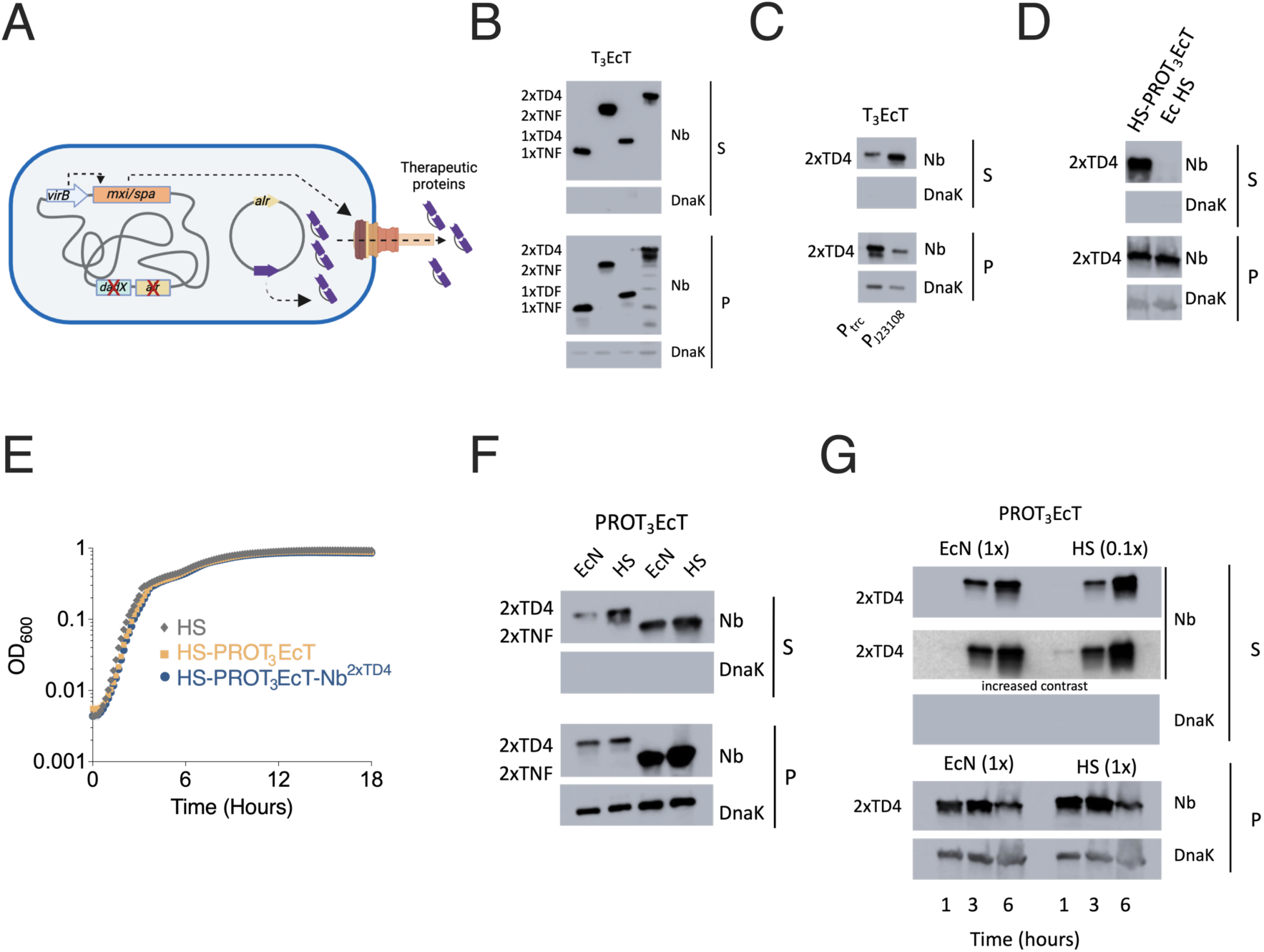
HS-PROT_3_EcT can be engineered to constitutively secrete SS^OspC2^-Nb^TD4^. (A) Schematic of HS-PROT3EcT. (B-D, F-G) Secretion assays of designated strains engineered to secrete noted FLAG-tagged nanobodies, each fused to an OspC2 secretion sequence. Supernatant (TCA precipitated) (S) and whole-cell pellet lysates (P) were obtained at 30 min (B), 1 hr (C), 3 hr (D, F), or at designated time points (G) post transfer of the bacteria to PBS (B) or LB (C-D, F-G). In the case of (B) and (C), IPTG was added to induce expression of the nanobodies. Immunoblots probed with anti-FLAG or anti-DnaK are shown. Blots are representative of three independent experiments. (E) Growth curves of HS *E. coli,* HS-PROT T_3_EcT, and HS-PROT_3_EcT-Nb^2x^^TD4^ grown in parallel. Data are representative of mean ± SEM of four technical repeats.

We next investigated the ability of HS-PROT_3_EcT to colonize the intestines of mice. C57BL/6 mice were orally inoculated twice with 10^8^ CFU of HS-PROT_3_EcT or diluent (PBS) (Fig. S5B). After reaching a stable level of colonization reflective of the shedding of ∼10^6^ CFU/g of feces (Fig. S5B), the mice were orally inoculated with food containing 10^8^ CFU of Cr(ΦStx2a). Mice colonized with HS-PROT_3_EcT, unlike those colonized with EcN-PROT_3_EcT, were as susceptible to Cr(ΦStx2a) infection as those solely pretreated with the diluent. They demonstrated no difference in the kinetics of expansion or level of Cr(ΦStx2a) colonization (Fig. 3D), weight loss (Fig. 3E), or survival (Fig. 3F). Thus, unlike EcN, *E. coli* HS does not appear to have inherent antibacterial properties, at least in protecting against infection with Cr. Thus, we focused on determining whether we could engineer this strain to prevent or delay the development of disease in mice infected with Cr(ΦStx2).

### HS-PROT_3_EcT efficiently secretes Nbs into its surroundings

We next investigated whether we could generate variants of Nb^TD4^ that were recognized as type III secreted proteins by outfitting them with a type III secretion sequence. Thus, we generated SS^OspC2^-Nb^1xTD4^ and SS^OspC2^-Nb^2xTD4^, monomeric and homodimeric Nb^TD4^, fused to the first 50 amino acids of OspC2, a native *Shigella* type III effector. These residues of OspC2 were previously established to support the secretion of a variety of Nbs^13,54^. As a comparator, we monitored the secretion of the previously characterized robustly secreted SS^OspC2^-Nb^1xTNF^ and SS^OspC2^-Nb^2xTNF^, monomeric and homodimeric anti-TNFα Nbs^13^. For these studies, we investigated the ability of each of these four nanobodies to be secreted by T_3_EcT, a variant of DH10b with the same constitutively active modified T3SS present in PROT_3_EcT^13^. SS^OspC2^-Nb^1xTD4^ and SS^OspC2^-Nb^2xTD4^ were secreted at levels similar to Nb^TNF^ (Fig. 4B).

For these initial studies, we characterized variants of Nb^TD4^ and Nb^TNF^ whose expression was under the control of an IPTG-inducible P*trc* promoter. However, as our goal was to develop variants of PROT_3_EcT that constitutively secrete Nb^TD4^ into the gut lumen, we next generated a variant of SS^OspC2^-Nb^2xTD4^ expressed under the control of the constitutive PJ23108 promoter on a plasmid that can be maintained via antibiotic or auxotrophic (*alr*) selection. Interestingly, in this case, we found that the constitutively expressed SS^OspC2^-Nb^2xTD4^ was more efficiently secreted from T_3_EcT (Fig. 4C).

Furthermore, SS^OspC2^-Nb^2xTD4^ was recognized as a secreted protein by HS-PROT_3_EcT but not HS *E. coli* (Fig. 4D), confirming that it was secreted in a type III secretion-dependent manner. In addition, we found that HS-PROT_3_EcT and HS-PROT_3_EcT- Nb^2xTD4^ exhibited essentially identical growth patterns as *E. coli* HS, indicating that the presence of an actively secreting modified type III secretion system does not result in a significant metabolic burden (Fig. 4E).

Next, to investigate whether HS-PROT_3_EcT secretes nanobodies at levels equivalent to EcN-PROT_3_EcT, we compared the secretory activities of two strains. We first assessed the ability of reach to secrete constitutively expressed SS^OspC2^-Nb^2xTNF^ and SS^OspC2^- Nb^2xTD4^. After a 3-hour incubation, we surprisingly found that HS-PROT_3_EcT secreted significantly higher levels of nanobodies, particularly in the case of SS^OspC2^-Nb^2xTD4^ (Fig. 4F). When we monitored secretion over a 6-hour time course, we observed that both strains continued to secrete the SS^OspC2^-Nb^2xTD4^ into their surroundings. In this case, to facilitate a parallel comparison of the secretion of Nb^2xTD4^ from both strains, we loaded 90% less of the supernatant fractions of HS-PROT_3_EcT. We again observed that HS- PROT_3_EcT more robustly secretes SS^OspC2^-Nb^2xTD4^. Future studies will address how the same secretion system in two different commensal *E. coli* differentially recognize secreted substrates.

### HS-PROT_3_EcT-Nb^2xTD4^ significantly delays the establishment of a Cr_Tir- M^EHEC^(ΦStx2a) infection

We next investigated whether pre-treatment with HS- PROT_3_EcT-Nb^2xTD4^ would protect mice from infection with Cr_Tir-M^EHEC^(ΦStx2a). Mice were colonized with HS-PROT_3_EcT-Nb^2xTD4^ (n=10) or HS-PROT_3_EcT or untreated inoculated with Cr_Tir-M^EHEC^(ΦStx2a) (Fig. 5A). As before (Fig. 3), mice were administered two doses of HS-PROT_3_EcT-Nb^2xTD4^ (n = 10) or HS-PROT_3_EcT (n = 10) or diluent (PBS) (n =5), this time separated by 6 days. On average, these mice shed both strains at ∼10^5^ CFU/g of feces (Fig. S5C). We observed some fluctuation in the levels of colonization of individual mice. While on occasion, HS-PROT_3_EcT CFU in the feces of a given mouse was below the limit of detection, only one mouse inoculated twice with HS- PROT_3_EcT-Nb^2xTD4^ never demonstrated evidence of colonization (Fig. S5C). This mouse was excluded from the study. Six days after their second PROT_3_EcT inoculation, the remaining 24 mice were inoculated with 1×10^8^ CFU of Cr_Tir-M^EHEC^(ΦStx2a) via the food inoculation model.

**Figure 5:**
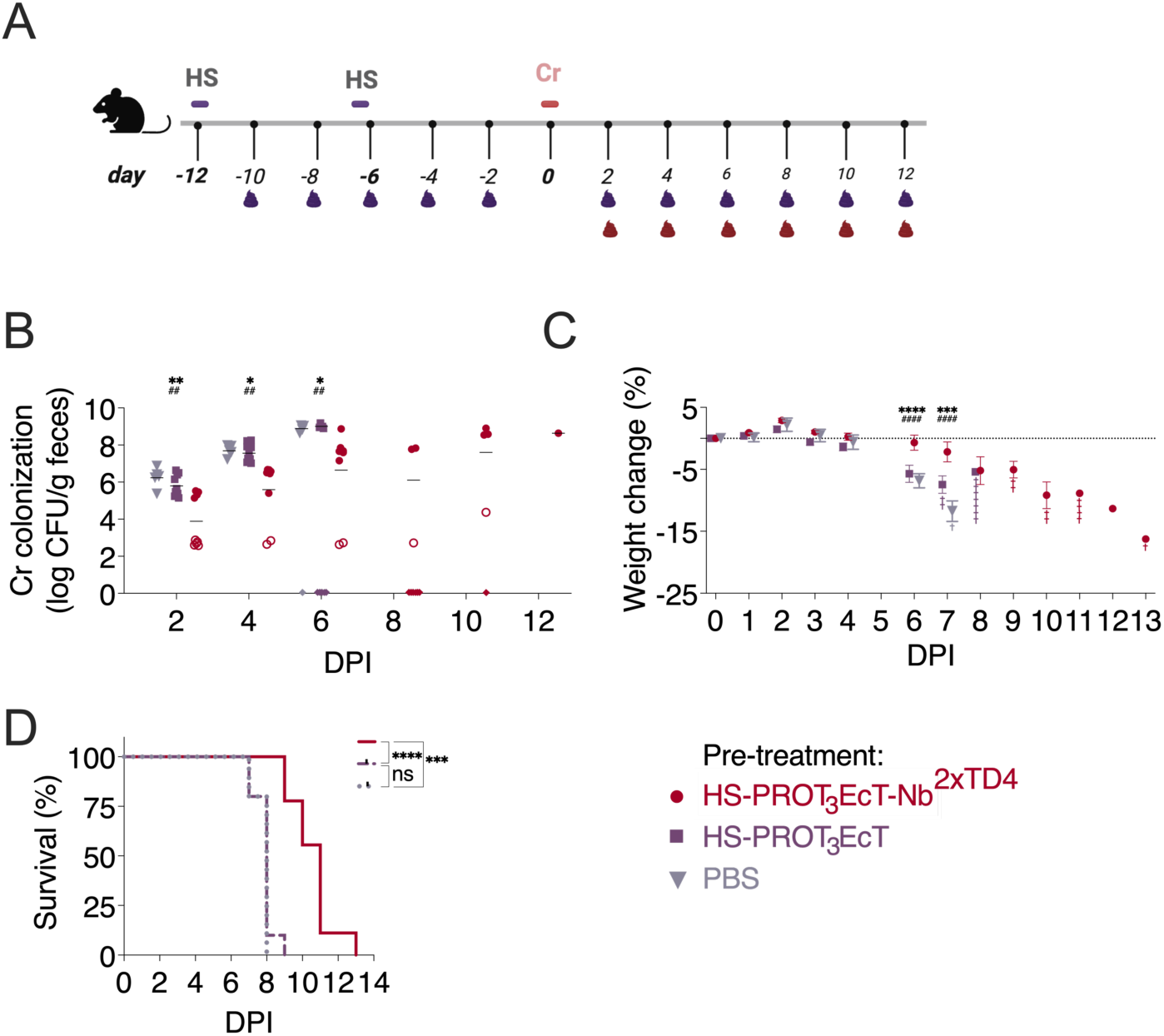
Pre-treatment with HS-PROT_3_EcT-TD4 delays Cr_Tir-M^EHEC^(ΦStx2a) colonization and prolongs survival of infected mice. (A) Study design schematic. (B-D) Seven-week-old female mice pre-treated with PBS (n=5), HS-PROT_3_EcT (n=10) or HS-PROT_3_EcT-Nb^2x^^TD4^ (N=9) were infected with 1×10^8^ CFU of Cr_Tir-M^EHEC^(ΦStx2a). (B) Viable counts of bacteria in feces were determined by plating. Each point shown represents an individual mouse, and each line represents the geometric mean. Shapes plotted on the x-axis indicate no data is available. Open symbols indicate CFU at the limit of detection (LOD). This value was used when calculating statistical significance. (C) Time course of body weight changes (%) over time. Mean +/-SEM plotted. Data in (B) and (C) were analyzed using two-way ANOVA with Bonferroni’s post hoc multiple comparison test at a 95% confidence interval. *Denotes comparison to PBS and ^#^ denotes comparison to HS-PROT_3_EcT. For (B), DPI 2 **p = 0.0035, ^##^p = 0.0040; DPI 4: *p = 0.0104, ^##^p = 0.0029; DPI 6: *p = 0.0114, ^##^p = 0.0034. For (C), DPI 6 ****p < 0.0001, ^####^p = 0.0001; DPI 7: ***p = 0.0007, ^####^p < 0.0001. (D) Kaplan–Meier survival curves of mice pretreated with HS-PROT_3_EcT-Nb^2x^^TD4^, HS-PROT_3_EcT, and Mock; infected with Cr_Tir-M^EHEC^(Stx2a). Statistical significance was determined by the log-rank (Mantel-Cox) test. All possible pairs of survival curves were compared independently. ***p = 0.0003, ****p < 0.0001, ns=non-significant.

Strikingly, compared to untreated mice or mice colonized with HS-PROT_3_EcT, those colonized with HS-PROT_3_EcT-Nb^2xTD4^ exhibited delayed Cr_Tir-M^EHEC^(ΦStx2a) colonization. On day 2 post-Cr_Tir-M^EHEC^(ΦStx2a) inoculation, Cr_Tir-M^EHEC^(ΦStx2a) levels were below the level of detection in 5/9 mice, two of which remained below the level of detection through day 6 (Fig. 5B). The kinetics of Cr_Tir-M^EHEC^(ΦStx2a) colonization, as measured by fecal shedding, was delayed in mice colonized by HS- PROT_3_EcT-Nb^2xTD4^ compared to mice pretreated with PBS or HS-PROT_3_EcT (Fig. 5B). These results are consistent with previous reports that Nb^1xTD4^ blocks pedestal formation *in vitro*^23^, a process that promotes Cr mucosal colonization^27^.

Consistent with the above results, compared to untreated mice or mice colonized with HS-PROT_3_EcT, mice colonized with HS-PROT_3_EcT-Nb^2xTD4^ exhibited delayed weight loss (Fig. 5C) and, on average, survived for 3.5 days longer (Fig. 5D). Delayed Cr_Tir- M^EHEC^(ΦStx2a) colonization generally correlated with prolonged survival.

To confirm that the secretion system remained functional in HS-PROT_3_EcT-Nb^2xTD4^ throughout the experiment, we used a plate secretion assay to evaluate the secretory activity of 10 of the last colonies isolated from each of the nine mice. We detected evidence of secreted nanobodies in all 90 colonies tested (Fig. S6), demonstrating that HS-PROT_3_EcT retains a functional secretion system within the intestines of mice for at least 25 days.

Given the published data that Nb^1xTD4^ prevents *in vitro* actin pedestal formation and that pedestal formation is essential for *C rodentium* gut colonization^27^, these results suggest that secreted Nb^2xTD4^ delays colonization when deposited into the intestinal lumen. As the Tir-M domain will not be accessible to luminal Nb^2xTD4^ until Tir is injected and inserted into the host cell membrane, these observations strongly suggest that colonization of HS-PROT_3_EcT-Nb^2xTD4^ results in the deposition of Nb^2xTD4^ in close proximity to intestinal epithelial cells, such that binds to and blocks Tir binding to Intiman. Whether HS-PROT_3_EcT-Nb^2xTD4^ establishes a replicative niche near the colonic epithelium remains to be determined.

## Summary

Here, to establish whether Nb^2xTD4^ can block or delay the onset of disease, we focused efforts on testing whether pre-colonization with HS-PROT_3_EcT-Nb^2xTD4^ blocks or delays infection with Shiga toxin-producing Cr. To enable these studies, we first developed an efficient, seamless gene replacement approach to generate a Cr strain that produced the Shiga toxin variant most closely associated with human disease and a chimeric Tir molecule that could be recognized by an Nb specific for EHEC Tir. While this protocol is similar to others^55^, it has the added advantage that all needed plasmids are generated via homologous or site-specific recombination rather than more time-consuming recombinant DNA techniques.

Next, interestingly, we found that EcN, but not *E. coli* HS, blocks infection with Cr, which led us to develop a variant of HS-PROT_3_EcT platform that can maintain each of its genetic components, including the operons encoding the modified T3SS, their shared transcriptional regulator and the therapeutic payload in the absence of antibiotic selection. Interestingly, HS-PROT_3_EcT outperformed EcN-PROT_3_EcT in its ability to secrete nanobodies outfitted with a type III secretion signal sequence, suggesting that there are differences in how the secreted substrates are recognized and delivered to the secretion apparatus with EcN and *E. coli* HS, an area of future investigation. With these modified strains in hand, we investigated whether the *in situ* secretion of Nb^2x^^TD4^ by HS- PROT_3_EcT inhibits the ability of Cr_Tir-M^EHEC^(ΦStx2a) to establish an infection. Remarkably, we found that the mice colonized with HS-PROT_3_EcT-Nb^2x^^TD4^ typically survived three more days than those colonized with HS-PROT_3_EcT.

Our long-term goal is to develop variants of PROT_3_EcT that can be used as an antibiotic-free means to treat intestinal-based infections. For this study, we focused on first determining whether the secreted anti-Tir Nb can provide prophylaxis against infection using Cr(ΦStx2a) as a model for EHEC. In future studies, given that EcN, as compared to *E. coli* HS, has inherent anti-Cr activity, we plan to test whether colonization with EcN- PROT_3_EcT- Nb^TD4^ completely blocks Cr(ΦStx2) from blocking an infection as well as if, when administered after the establishment of a Cr(ΦStx2a), such that it can be used as a treatment modality. We will also investigate whether variants of PROT_3_EcT outfitted to deliver a cocktail of therapeutic payloads, secreted nanobodies that block the activity of additional CR(ΦStx2)/EHEC virulence factors, i.e., intimin and Stx2^56,57^, provide enhanced protection in both prophylaxis and disease models. As it is thought that treatment with antibiotics increases the risk of development of HUS due to increased production and secretion of Stx2, we also plan to investigate whether co-treatment with the PROT_3_EcT that secrete anti-Tir and anti-Stx2 Nbs can protect from developing HUS when antibiotics are used as a treatment modality.

While this study was limited to investigating the ability of a secreted anti-Tir Nb to inhibit the establishment of an intestinal infection, we previously established that a secreted anti-TNFα Nb blocked the development of colitis in a mouse pre-clinical model of inflammatory bowel disease^13^. Furthermore, we have demonstrated that PROT_3_EcT can be engineered to secrete Nbs and other heterologous proteins fused to an N-terminal secretion sequence. Together, these complementary studies illustrate the power and versatility of the PROT_3_EcT platform in terms of *E. coli* chassis, therapeutic payload, and disease applications and suggest that this platform can be extended to treat additional gut-based diseases as well as those linked to the gut microbiota.

## Materials and Methods

Plasmids and strains are summarized in Tables S1 and S2, while Sequences of oligos and synthetic DNA fragments are cataloged in Tables S3 and S4.

### Bacterial growth conditions

Unless otherwise noted, *E. coli* and Cr strains were grown in Luria broth (LB: 10g/L tryptone, 5g/L yeast extract, 10g/L NaCl) or minimal media (1xM9 salts, 0.5% glycerol, 2 mM MgSO_4_, 0.1mm CaCl_2_, 1 *μ*g/ml thiamine, 1 *μ*g/ml biotin) at 37°C with aeration on a roller or on solid media (15% agar). Strains transformed with temperature-sensitive plasmid pCP20 or pTKRED were maintained at 30°C and cured by incubation at 42°C. When noted, antibiotics (100 *μ*g/ml spectinomycin, 100 *μ*g/ml ampicillin, 50 *μ*g/ml kanamycin, 12.5 *μ*g/ml tetracycline,10 *μ*g/ml chloramphenicol, 100 *μ*g/ml hygromycin), D-alanine (50 *μ*g/ml),1mM IPTG (1 mM) were added. Fusaric acid solid media plates were made as described in ^58^.

### Plasmid construction

#### Donor plasmids

The Donor plasmids were generated via the gateway recombination system. Synthetic DNA sequence fragments [attB1-Stx hybrid-attB2], [attB1- EHEC_Cr_Tir-M-attB2], and [attB1-EPEC_Cr_Tir-M-attB2] (Twist Bioscience) were introduced into pDONR221 via BP reactions Gateway.

#### Nb^TD4^ expression plasmids

The Ptac-regulated SS^OspC2^Nb^TD4^ expression plasmids were generated via the gateway recombination system^32^. Synthetic DNA sequences [attb1-SS^OspC2^-Nb^1x^^TD4^-attB2] and [attb1-SS^OspC2^-Nb^2x^^TD4^-attB2] (Twist Bioscience) were introduced into pDONR221 via Gateway BP reactions. Subsequently, [attb1-SS^OspC2^-Nb ^1x^^TD4^-attB2] and [attb1-SS^OspC2^-Nb ^2x^^TD4^-attB2] were introduced into pDSW206-ccdB-FLAG via LR reactions to create pDSW206-SS^OspC2^-Nb^1x^^TDF^ and pDSW206-SS^OspC2^- Nb^2x^^TDF^. The constitutively expressed SS^OspC2^-Nb^2x^^TDF^ was introduced via restriction/ligation cloning using a synthetic DNA fragment [EcoRI-PJ23108-SS^OspC2^-Nb ^2x^^TD4^-XbaI], which encodes the Nb with a synthetic 5’ untranslated region (UTR) and the constitutive PJ23108 into pCPG-alr. Both were digested with *EcoRI/XbaI*. The insert was sequence verified.

### Strain construction

#### Cr(<ΙStx2a)

*<u>Step 1</u>*: A fragment of DNA composed of a Tet^R^ cassette flanked by I-SceI

#### Cr(ΦStx2a)

*<u>Step 1</u>*: A fragment of DNA composed of a Tet^R^ cassette flanked by I-SceI sites and 427/408 base pairs of homology up/down-stream of *stx2dact* in Cr(ΦStx2dact) ([Stx^UP^-SceI-TetR-SceI-Stx^DN^]) was generated by 3-piece splicing by overlap-extension (SOEing) PCR. The upstream and downstream regions of homology were PCR amplified from Cr(ΦStx2dact) using P1/P2 and P3/P4. The center SceI-Tet^R^-SceI fragment was amplified from *E. coli* atg/gid::landing pad^54^ using P5/P6. The three pieces were combined using P1/P4 primers. *<u>Step 2</u>*: Cr(ΦStx2dact) containing pTKRED was transformed with [Stx^UP^-SceI-TetR-SceI-Stx^DN^] and λ-red recombineering technology^33,34^ was used to generate Cr(ΦStx2::Tet^R^). pTRKRED was cured, and the insert was PCR verified using P6/P7. *<u>Step 3</u>*: Cr(ΦStx2::Tet^R^) was transformed with pTKRED and pDonor-Stx2A. *<u>Step 4</u>*: An overnight culture of Cr(ΦStx2::Tet^R^) plus the plasmids was back-diluted 1:100 into M9 media/spectinomycin and grown at 30°C. After 2 h, 1 mM IPTG was added. The culture was incubated for an additional 8 h, at which point 0.3% arabinose and 1 mM IPTG were added, and the culture was incubated o/n at 30°C. The following day, 50 ul of the culture was spread on an M9 + FA plate and incubated o/n at 37°C. The next day, individual colonies were patched onto an LB plate and were subsequently screened for those that were Tet^S^/Spec^S^/Kan^S^. <u>Step 5</u>: The resulting strain, Cr(ΦStx2a), was verified by PCR using primers P8/P9, and the amplified fragment containing the swapped sequence was sequence verified.

#### Cr_Tir-M^EPEC^(ΦStx2dact), Cr_Tir-M^EHEC^(ΦStx2dact), Cr_Tir-M^EPEC^(ΦStx2a) and Cr_Tir-M^EHEC^(ΦStx2a)

*<u>Step 1</u>*: A fragment of DNA composed of a Tet^R^ cassette flanked by I-SceI sites and 300 basepairs of homology up-/downstream of Cr *tir-M* was generated by 3-piece SOEing PCR ([Tir^UP^-SceI-TetR-SceI-Tir^DN^]). The upstream and downstream regions of homology were PCR amplified from Cr(ΦStx2dact) using P10/P11 and P12/P13. The middle SceI-Tet^R^-SceI fragment was amplified from *E. coli* atg/gid::landing pad using P5/P6. The three pieces were combined using P10/P13 primers. *<u>Step 2</u>*: Cr(ΦStx2dact) and Cr(ΦStx2a) carrying pTKRED were transformed with Tir^UP^-SceI-TetR-SceI-Tir^DN^ and 1-Red recombineering was used to generate Cr-Tir- M^TetR^(ΦStx2dact) and Cr-Tir-M^TetR^(ΦStx2a). pTRKRED was cured, and the inserts were PCR verified using P6/P14. *<u>Step 3</u>*: Cr(ΦStx2::Tet^R^) and Cr-Tir-M^TetR^(ΦStx2a) were each retransformed with pTKRED plus pDonor-Cr-Tir-M^EHEC^ or pDonor-Cr-Tir-M^EPEC^. *<u>Step 4</u>*: as above for generating Cr(ΦStx2a). *<u>Step 5</u>*: The inserts in the final strains, Cr_Tir- M^EPEC^(ΦStx2dact), Cr_Tir-M^EHEC^(ΦStx2dact), Cr_Tir-M^EPEC^(ΦStx2a) and Cr_Tir- M^EHEC^(ΦStx2a), were verified by PCR using primers P15/P16 and the amplified fragments contained the swap regions of DNA were sequence verified.

#### HS-PROT_3_EcT

*<u>Step 1</u>*: A synthetic 1.3 landing pad insertion site (LP1-TetR-LP2) was introduced into the *yieN/trkB* locus of PROT_3_EcT-2. The landing pad fragment was PCR amplified from PROT3EcT-1- LP^yie/trk^ using P17/18 primers. This DNA was introduced into PROT_3_EcT-2 containing pTKRED, and λ-Red recombineering was used to generate PROT_3_EcT-2-LP^yie/trk^. pTKRED was cured. *<u>Step 2</u>*: PROT_3_EcT-2-LP^yie/trk^ was transformed with pTKRED and pTKIP-PJ23119-virB, and the landing pad recombination system^34^ was used to introduce the *virB* expression cassette via hygromycin selection into its chromosome at the yie/trk locus to generate PROT3EcT-2-virB-hygro. Integration was confirmed by PCR with P17/P19 and P18/P20. <u>Step 3</u>: The KAN^R^ and Hygro^R^ FRT cassettes in PROT3EcT-2-virB-hygro were removed using the FLP recombinase (pCP20) to generate PROT3EcT-2-virB. *<u>Step 4</u>*: The lambda red recombination system^31,32^ was used to sequentially delete dadX and alr from PROT3EcT-2-virB using oligomers (P21/P22 and P23/P24), respectively. The KAN^R^ was removed from the dadX locus before proceeding to delete *alr*. Deletions were confirmed by PCR with oligomers (P25/P26 and P27/P28, respectively). When both *alr* and *dadX* were removed, the strain was grown in the presence of D-ala. The final strain is referred to as HS-PROT_3_EcT.

#### Bacterial growth curves

Overnight bacteria cultures were back-diluted (1:40) in LB. 200ul of each culture was placed in quadruplicate into a 96-well plate (Corning). A Breathe-Easy film (Sigma-Aldrich) was applied to minimize evaporation. The plate was incubated at 37°C with shaking, and OD_600_ readings were obtained every ten minutes using a SpectraMax i3x Plate Reader (Molecular Devices). Growth curves were plotted using GraphPad Prism version 10 (GraphPad Software, Inc., San Diego, CA, USA).

#### Mouse infection studies

Six-week-old female C57BL/6J mice purchased from Jackson Labs were used for all experiments. Upon arrival, they were given at least one week to acclimate. Mice were housed in microisolator cages under specific pathogen-free conditions in the barrier facility at Tufts University School of Medicine. Five mice were randomly grouped in each cage. Mice received bacteria via mouth pipetting, oral gavage, or food inoculation. Before receiving Cr, food was held the night for 12 h. After inoculation with the *E coli* HS- or EcN-based strains, mice were weighed every 1-2 days. After receiving Cr, the mice were weighed and observed for clinical signs of disease each day. Mice with greater than 15% body weight loss, with or without signs of distress, were sacrificed by CO_2_ inhalation followed by cervical dislocation.

#### Fecal shedding assay

Fecal pellets were collected and weighed. The pellets were homogenized in 200 ul of PBS by mashing using wide-mouth pipette tips. After which, they were serially diluted and plated on LB agar plates with ampicillin for detecting EcN- PROT_3_EcT, kanamycin for detecting HS-PROT_3_EcT, and chloramphenicol for detecting Cr. The next day, colonies were enumerated, and the total CFU was calculated.

#### Tir secretion assay

Cultures inoculated with single colonies of Cr(ΦStx2dact), Cr_Tir- M^EHEC^(ΦStx2a) or Cr_Tir-M^EPEC^(ΦStx2a) were incubated o/n with aeration at 37°C. In the AM, the cultures were back-diluted 1:50 into 5 ml of DMEM/0.1M HEPES and incubated without aeration at 37°C in a 5% CO2 incubator. After 6 h, 2 ml of each culture was centrifuged twice at 12000 rpm for 10 minutes. Proteins in the supernatants were precipitated with 10% (v/v) trichloroacetic acid (TCA). The supernatant/pellet fractions were resuspended in 50/100 μL of protein loading dye. 10 ul of each fraction were loaded onto a 12% Tris-Glycine gel (Novex), which was transferred to a nitrocellulose membrane and blotted with an anti-Tir antibody^19^ and anti-GroEL (1:100,000) antibody (Abcam ab69617). GroEL was used as a loading and lysis control.

#### *E. coli* liquid secretion assays

Liquid secretion assays were performed as previously described^13^ with some modifications. Overnight cultures of *E. coli* grown in LB at 37°C were back diluted 1:50. For assays requiring IPTG induction, 1mM IPTG was added to each culture at the start of the back dilution. Once cultures reached OD_600_ of 1.2-1.5, the bacteria were pelleted and resuspended in 2.5 mL of fresh LB or PBS, as noted, and incubated at 37°C on a roller. After designated periods of time, total cell and supernatant fractions were separated by centrifugation at 13,000 rpm for 1 min. The cell pellet was taken as the whole cell lysate fraction. The supernatant fraction was subjected to a second centrifugation step to remove any remaining bacteria. For each set of experiments, the volume of bacteria centrifuged was normalized to the OD_600_ reading of the slowest growing culture to account for differences in bacterial titers. Samples were not normalized for the time course assays. The pellet was resuspended in 100 uL. Proteins in the supernatant were precipitated with trichloroacetic acid (TCA) (10% v/v) and resuspended in 50 uL. Proteins resuspended in loading dye were incubated at 95°C for 10 min. Ten microliters of TCA-precipitated supernatant samples (20%) and five microliters of the pellet (5%) were loaded onto a 12% Tris-Glycine SDS-PAGE gel for analysis (i.e., the ratio of supernatant to pellet samples loaded was 3:2). Proteins were transferred to nitrocellulose membranes and immunoblotted with mouse anti-M2-FLAG (1:5,000) (Sigma) or mouse anti-DnaK (1:5,000) (ab69617).

#### Solid plate secretion assay

Solid plate secretion assays were performed as previously described^59^. Briefly, single colonies grown overnight in 96-well plates were quad-spotted onto a solid agar using a using a BMC3-BC pinning robot (S&P Robotics). A 384-pin tool was used the following day to transfer equivalent amounts of bacteria to a solid media-containing plate over which a nitrocellulose membrane was laid immediately. All incubations were carried out at 37°C. After 6 h, the membrane was removed, washed to remove adherent bacteria, and immunoblotted with an anti-M2-FLAG antibody (1:5000, Sigma) to detect the secreted epitope-tagged nanobodies.

#### Statistical analyses

Statistical analyses were performed using GraphPad Prism software (version 10). Specific tests used are indicated in the figure legends. Significant difference is indicated as *p<0.05, **p <0.01, ***p<0.001, ****p<0.0001, and ns = non-significant for all figures.

## Acknowledgments

We thank Dr. Joan Mecsas for reading the paper and advising on statistical analyses and members of the Lesser, Leong, Barczak and Goldberg labs and the Tufts CMS for helpful discussions and advice. Some of the graphics were created with BioRender.com. This work was supported by NIH DK113599, the Brit d’Arbeloff Research Scholar award to C.F.L, and a Massachusetts General Hospital Executive Committee on Research Fund for Medical Discovery Postdoctoral Fellowship Award to C. G.-P.

## Data availability

All experimental data described in this study are included in the manuscript and/or supporting information.

## References

1. Cubillos-Ruiz, A. et al. Engineering living therapeutics with synthetic biology. Nat Rev Drug Discov 20, 941–960 (2021).

2. McNerney, M. P., Doiron, K. E., Ng, T. L., Chang, T. Z. & Silver, P. A. Theranostic cellsti emerging clinical applications of synthetic biology. Nat Rev Genet 22, 730–746 (2021).

3. Lynch, J. P., Goers, L. & Lesser, C. F. Emerging strategies for engineering *Escherichia coli* Nissle 1917-based therapeutics. Trends Pharmacol Sci 43, 772–786 (2022).

4. Russell, B. J. et al. Intestinal transgene delivery with native *E. coli* chassis allows persistent physiological changes. Cell 185, 3263–3277.e15 (2022).

5. Lodinová-Zádniková, R. & Sonnenborn, U. Effect of preventive administration of a nonpathogenic *Escherichia coli* strain on the colonization of the intestine with microbial pathogens in newborn infants. Biol Neonate 71, 224–232 (1997).

6. Din, M. O. et al. Synchronized cycles of bacterial lysis for *in vivo* delivery. Nature 536, 81–85 (2016).

7. Gurbatri, C. R., Arpaia, N. & Danino, T. Engineering bacteria as interactive cancer therapies. Science 378, 858–864 (2022).

8. Praveschotinunt, P., et al. Engineered *E. coli* Nissle 1917 for the delivery of matrix- tethered therapeutic domains to the gut. Nat Commun 10, 5580 (2019).

9. Piñero-Lambea, C. et al. Programming controlled adhesion of *E. coli* to target surfaces, cells, and tumors with synthetic adhesins. ACS Synth Biol 4, 463–473 (2015).

10. Deng, W. et al. Assembly, structure, function and regulation of type III secretion systems. Nat Rev Microbiol 15, 323–337 (2017).

11. Worrall, L. J., Majewski, D. D. & Strynadka, N. C. J. Structural Insights into Type III Secretion Systems of the Bacterial Flagellum and Injectisome. Annu Rev Microbiol 77, 669–698 (2023).

12. Ménard, R., Sansoneg, P. J. & Parsot, C. Nonpolar mutagenesis of the ipa genes defines IpaB, IpaC, and IpaD as effectors of *Shigella flexner*i entry into epithelial cells. J Bacteriol 175, 5899–5906 (1993).

13. Lynch, J. P. et al. Engineered *Escherichia coli* for the in situ secretion of therapeutic nanobodies in the gut. Cell Host & Microbe 31, 634–649.e8 (2023).

14. González-Prieto, C., Lynch, J. P. & Lesser, C. F. PROT3EcT, engineered *Escherichia coli* for the targeted delivery of therapeutics. Trends Mol Med S1471-4914(23)00157–0 (2023) doi:10.1016/j.molmed.2023.07.007.

15. Tobe, T. et al. An extensive repertoire of type III secretion effectors in *Escherichia coli* O157 and the role of lambdoid phages in their dissemination. Proc Natl Acad Sci U S A 103, 14941–14946 (2006).

16. de Grado, M., et al. Identification of the intimin-binding domain of Tir of enteropathogenic *Escherichia coli*. Cell Microbiol 1, 7–17 (1999).

17. Hartland, E. L., et al. Binding of intimin from enteropathogenic *Escherichia coli* to Tir and to host cells. Mol Microbiol 32, 151–158 (1999).

18. Kenny, B., et al. Enteropathogenic E. coli (EPEC) transfers its receptor for intimate adherence into mammalian cells. Cell 91, 511–520 (1997).

19. Mallick, E. M. et al. The ability of an akaching and effacing pathogen to trigger localized actin assembly contributes to virulence by promoting mucosal akachment. Cell Microbiol 16, 1405–1424 (2014).

20. Melton-Celsa, A. R. Shiga Toxin (Stx) Classification, Structure, and Function. Microbiol Spectr 2, 2.4.06 (2014).

21. Wen, S. X., Teel, L. D., Judge, N. A. & O’Brien, A. D. A plant-based oral vaccine to protect against systemic intoxication by Shiga toxin type 2. Proc Natl Acad Sci U S A 103, 7082–7087 (2006).

22. McGannon, C. M., Fuller, C. A. & Weiss, A. A. Different classes of antibiotics differentially influence shiga toxin production. AnHmicrob Agents Chemother 54, 3790–3798 (2010).

23. Ruano-Gallego, D., et al. A nanobody targeting the translocated intimin receptor inhibits the akachment of enterohemorrhagic *E. coli* to human colonic mucosa. PLoS Pathog 15, e1008031 (2019).

24. Mohawk, K. L. & O’Brien, A. D. Mouse Models of *Escherichia coli* O157:H7 Infection and Shiga Toxin Injection. J Biomed Biotechnol 2011, 258185 (2011).

25. Eaton, K. A., Fontaine, C., Friedman, D. I., Conti, N. & Alteri, C. J. Pathogenesis of Colitis in Germ-Free Mice Infected With EHEC O157:H7. Vet Pathol 54, 710–719 (2017).

26. Mallick, E. M., et al. Allele- and tir-independent functions of intimin in diverse animal infection models. Front Microbiol 3, 11 (2012).

27. Deng, W., Vallance, B. A., Li, Y., Puente, J. L. & Finlay, B. B. *Citrobacter rodenHum* translocated intimin receptor (Tir) is an essential virulence factor needed for actin condensation, intestinal colonization and colonic hyperplasia in mice. Molecular Microbiology 48, 95–115 (2003).

28. Mallick, E. M. et al. A novel murine infection model for Shiga toxin–producing *Escherichia coli*. J. Clin. Invest. 122, 4012–4024 (2012).

29. Luperchio, S. A. & Schauer, D. B. Molecular pathogenesis of *Citrobacter rodenHum* and transmissible murine colonic hyperplasia. Microbes Infect 3, 333–340 (2001).

30. Thorpe, C. M., Pulsifer, A. R., Osburne, M. S., Vanaja, S. K. & Leong, J. M. *Citrobacter rodenHum*(ϕStx2dact), a murine infection model for enterohemorrhagic *Escherichia coli*. Current Opinion in Microbiology 65, 183–190 (2022).

31. Flowers, L. J., Bou Ghanem, E. N. & Leong, J. M. Synchronous Disease Kinetics in a Murine Model for Enterohemorrhagic *E. col*i Infection Using Food-Borne Inoculation. Front. Cell. Infect. Microbiol. 6, (2016).

32. Marsischky, G. & LaBaer, J. Many Paths to Many Clones: A Comparative Look at High- Throughput Cloning Methods. Genome Res. 14, 2020–2028 (2004).

33. Datsenko, K. A. & Wanner, B. L. One-step inactivation of chromosomal genes in Escherichia coli K-12 using PCR products. Proc. Natl. Acad. Sci. U.S.A. 97, 6640–6645 (2000).

34. Kuhlman, T. E. & Cox, E. C. Site-specific chromosomal integration of large synthetic constructs. Nucleic Acids Res 38, e92 (2010).

35. Ogura, Y. et al. The Shiga toxin 2 production level in enterohemorrhagic *Escherichia coli* O157:H7 is correlated with the subtypes of toxin-encoding phage. Sci Rep 5, 16663 (2015).

36. Deng, W. et al. Dissecting virulence: systematic and functional analyses of a pathogenicity island. Proc Natl Acad Sci U S A 101, 3597–3602 (2004).

37. Mills, E., Baruch, K., Charpentier, X., Kobi, S. & Rosenshine, I. Real-time analysis of effector translocation by the type III secretion system of enteropathogenic *Escherichia coli*. Cell Host Microbe 3, 104–113 (2008).

38. Lai, Y., Rosenshine, I., Leong, J. M. & Frankel, G. Intimate host akachment: enteropathogenic and enterohaemorrhagic Escherichia coli. Cell Microbiol 15, 1796–1808 (2013).

39. Kenny, B. Phosphorylation of tyrosine 474 of the enteropathogenic *Escherichia coli* (EPEC) Tir receptor molecule is essential for actin nucleating activity and is preceded by additional host modifications. Mol Microbiol 31, 1229–1241 (1999).

40. DeVinney, R. et al. Enterohemorrhagic *Escherichia coli* O157:H7 Produces Tir, Which Is Translocated to the Host Cell Membrane but Is Not Tyrosine Phosphorylated. Infect Immun 67, 2389–2398 (1999).

41. Campellone, K. G., Robbins, D. & Leong, J. M. EspFU is a translocated EHEC effector that interacts with Tir and N-WASP and promotes Nck-independent actin assembly. Dev Cell 7, 217–228 (2004).

42. Garmendia, J. et al. TccP is an enterohaemorrhagic *Escherichia coli* O157:H7 type III effector protein that couples Tir to the actin-cytoskeleton. Cell Microbiol 6, 1167– 1183 (2004).

43. Kenny, B. The enterohaemorrhagic *Escherichia coli* (serotype O157:H7) Tir molecule is not functionally interchangeable for its enteropathogenic E. coli (serotype O127:H6) homologue. Cell Microbiol 3, 499–510 (2001).

44. DeVinney, R., Puente, J. L., Gauthier, A., Goosney, D. & Finlay, B. B. Enterohaemorrhagic and enteropathogenic *Escherich*ia coli use a different Tir-based mechanism for pedestal formation. Mol Microbiol 41, 1445–1458 (2001).

45. Mallick, E. M., et al. Allele- and Tir-Independent Functions of Intimin in Diverse Animal Infection Models. FronHers in Microbiology 3, (2012).

46. Likle, D. J. & Coombes, B. K. Molecular basis for CesT recognition of type III secretion effectors in enteropathogenic *Escherichia coli*. PLoS Pathog 14, e1007224 (2018).

47. Sonnenborn, U. & Schulze, J. The non-pathogenic *Escherichia coli* strain Nissle 1917 – features of a versatile probiotic. Microbial Ecology in Health and Disease 21, 122– 158 (2009).

48. Deriu, E. et al. Probiotic bacteria reduce *salmonella typhimuriu*m intestinal colonization by competing for iron. Cell Host Microbe 14, 26–37 (2013).

49. Sassone-Corsi, M. et al. Microcins mediate competition among Enterobacteriaceae in the inflamed gut. Nature 540, 280–283 (2016).

50. Leatham, M. P. et al. Precolonized Human Commensal *Escherichia coli* Strains Serve as a Barrier to E. coli O157:H7 Growth in the Streptomycin-Treated Mouse Intestine. Infect Immun 77, 2876–2886 (2009).

51. Levine, M. M., et al. *Escherichia coli* strains that cause diarrhoea but do not produce heat-labile or heat-stable enterotoxins and are non-invasive. Lancet 1, 1119–1122 (1978).

52. Imai, J., et al. Flagellin-mediated activation of IL-33-ST2 signaling by a pathobiont promotes intestinal fibrosis. Mucosal Immunol 12, 632–643 (2019).

53. Matsumoto, M. et al. Free D-amino acids produced by commensal bacteria in the colonic lumen. Sci Rep 8, 17915 (2018).

54. Reeves, A. Z. et al. Engineering *Escherichia coli* into a protein delivery system for mammalian cells. ACS Synth Biol 4, 644–654 (2015).

55. Yang, J. et al. High-efficiency scarless genetic modification in *Escherichia col*i by using lambda red recombination and I-SceI cleavage. Appl Environ Microbiol 80, 3826– 3834 (2014).

56. Tremblay, J. M. et al. A Single VHH-Based Toxin-Neutralizing Agent and an Effector Antibody Protect Mice against Challenge with Shiga Toxins 1 and 2. Infect Immun 81, 4592–4603 (2013).

57. Gelfat, I. et al. Single domain antibodies against enteric pathogen virulence factors are active as curli fiber fusions on probiotic *E. coli Nissle* 1917. PLoS Pathog 18, e1010713 (2022).

58. Bochner, B. R., Huang, H. C., Schieven, G. L. & Ames, B. N. Positive selection for loss of tetracycline resistance. J Bacteriol 143, 926–933 (1980).

59. Ernst, N. H., Reeves, A. Z., Ramseyer, J. E. & Lesser, C. F. High-Throughput Screening of Type III Secretion Determinants Reveals a Major Chaperone-Independent Pathway. mBio 9, e01050–18 (2018).

