## Supplemental Materials for "*In situ* deposition of nanobodies by an engineered commensal microbe promotes survival in a mouse model of enterohemorrhagic *E. coli*"

**This PDF file includes:**

Figures S1 to S6
Tables S1 to S4
SI References

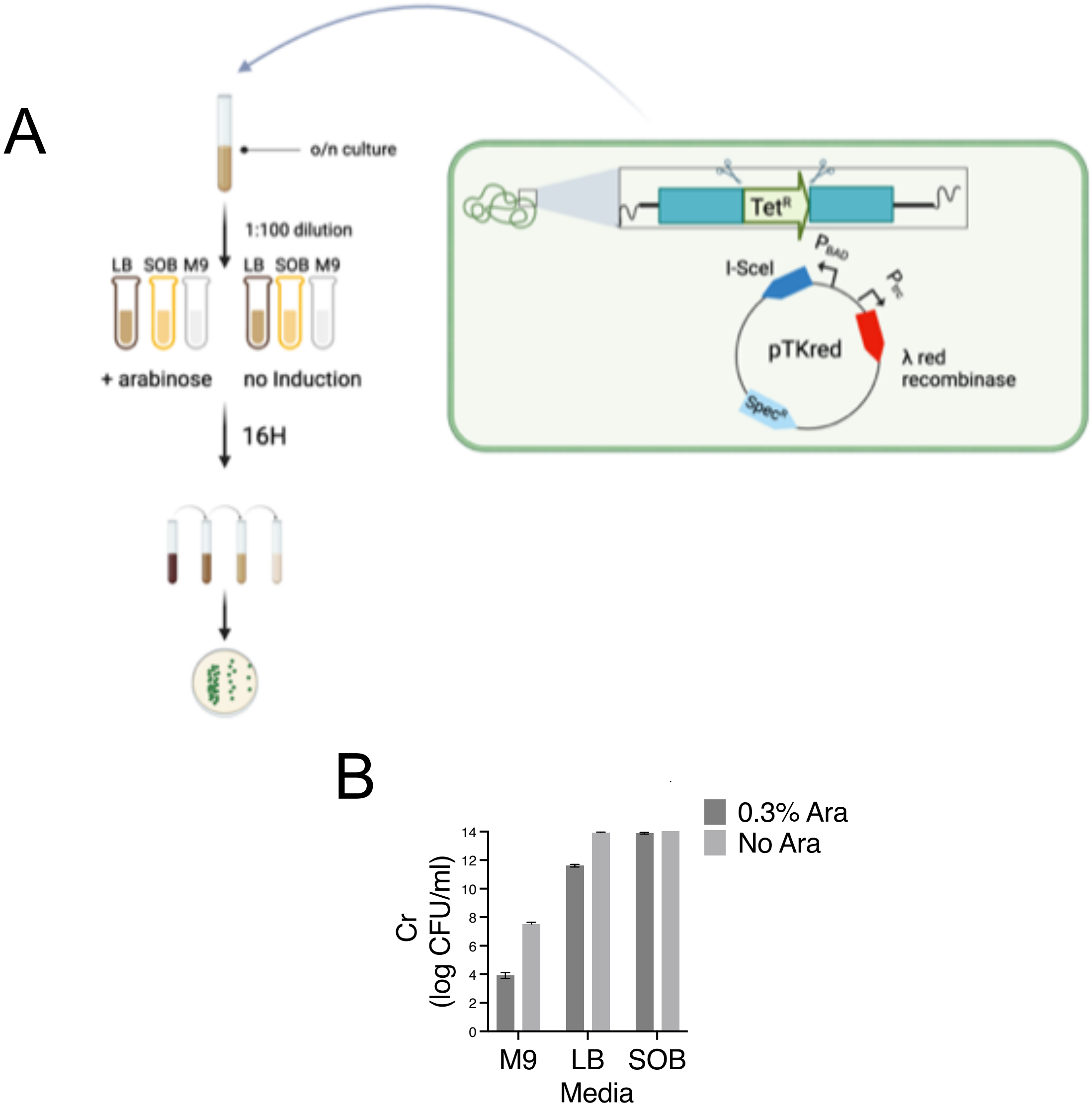


**Fig S1: Comparison of arabinose induction efficiency in minimal vs. rich media**

(A) Experimental workflow to determine the efficiency of arabinose-induced expression of I-SceI in minimal vs. rich media. Overnight cultures of Cr engineered with a chromosomally integrated $Tet^R$ cassette flanked by SceI sites [Cr(ΦStx2::$Tet^R$)] that carry pTKRED, a temperature-sensitive plasmid that expresses I-SceI under the control of $P_{BAD}$, an arabinose-inducible promoter, were back-diluted into LB, SOB or M9 minimal media in the presence and absence of 0.3% arabinose and incubated overnight (16 h) at 30ºC. Serial cultures were subsequently plated on LB media, and CFUs were enumerated. (B) Quantification of viable bacterial CFU recovered from the six designated conditions. The data shown is representative of three technical repeats, which were repeated at least three times.

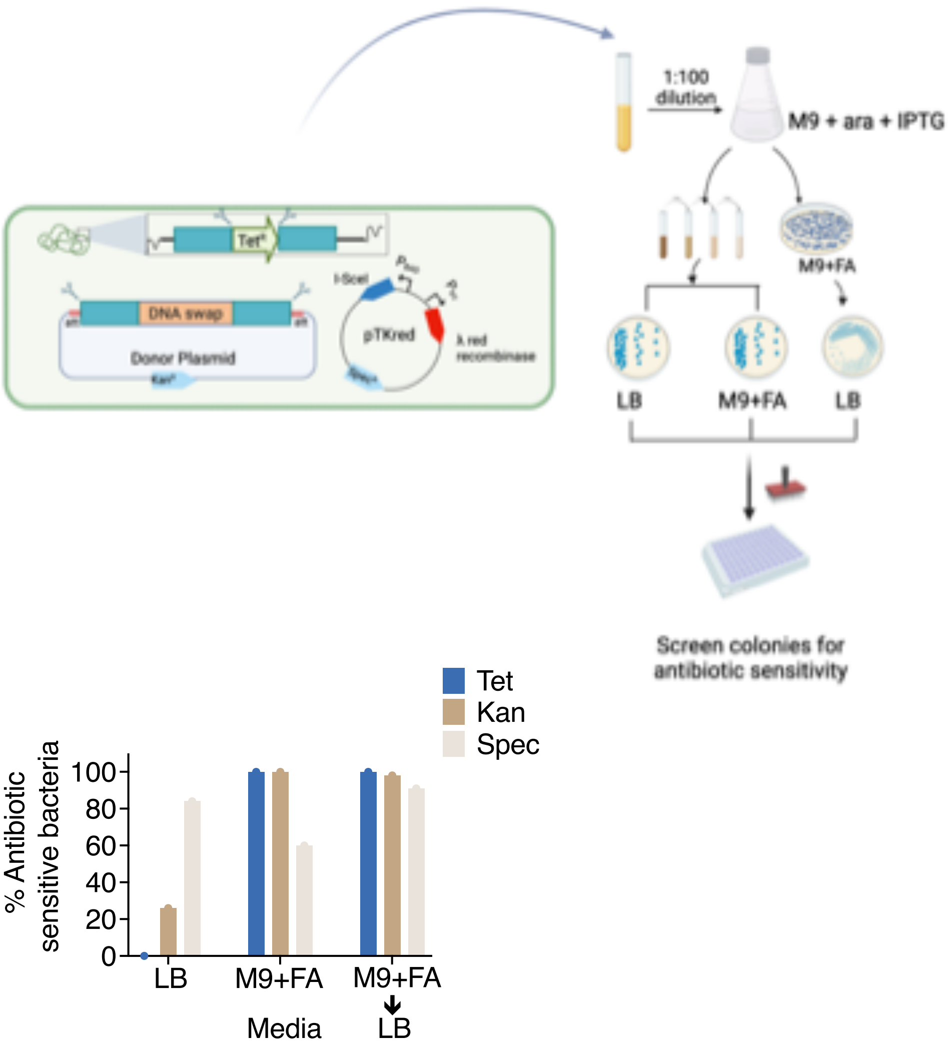


**Fig S2: Fusaric acid counter selection enhances the selection of Tet$^S$ colonies.** (A) Experimental workflow to identify conditions that improve the isolation for Cr variants engineered with seamless DNA swaps. Overnight cultures of Cr engineered with a chromosomally integrated Tet$^R$ cassette flanked by SceI sites [Cr(ΦStx2::Tet$^R$)] that carry pTKRED, and a donor plasmid (pDonor-Stx2A) were back-diluted into M9 minimal media + 0.3% arabinose + 1 mM IPTG and incubated overnight (16h). In the AM, serial dilutions of the cultures were plated onto LB and M9 + FA. In parallel, the undiluted culture was directly plated on M9+FA media and then streaked out for single colonies on LB plates. Forty-five single isolates obtained from each condition were screened for antibiotic sensitivity. (B) The percentage of antibiotic-sensitive bacteria recovered from each experimental cohort. The data shown is representative of three technical repeats, which were repeated at least three times.

```
Stx2a_A         MKCILFKWVLCLLLGFSSVSYSREFTIDFSTQQSYVSSLNSIRTEISTPLEHISQGTTSV    60
Stx2dact_A      MKCILFKWVLCLLLGFSSVSYSREFMIDFSTQQSYVSSLNSIRTEISTPLEHISQGTTSV    60
                ************************* **********************************

Stx2a_A         SVINHTPPGSYFAVDIRGLDVYQARFDHLRLIIEQNNLYVAGFVNTATNTFYRFSDFTHI    120
Stx2dact_A      SVINHTPPGSYFAVDIRGLDVYQARFDHLRLIIEQNNLYVAGFVNTATNTFYRFSDFTHI    120
                ************************************************************

Stx2a_A         SVPGVTTVSMTTDSSYTTLQRVAALERSGMQISRHSLVSSYLALMEFSGNTMTRDASRAV    180
Stx2dact_A      SVPGVTTVSMTTDSSYTTLQRVAALERSGMQISRHSLVSSYLALMEFSGNTMTRDASRAV    180
                ************************************************************

Stx2a_A         LRFVTVTAEALRFRQIQREFRQALSETAPVYTMTPGDVDLTLNWGRISNVLPEYRGEDGV    240
Stx2dact_A      LRFVTVTAEALRFRQIQREFRQALSETAPVYTMTPEEVDLTLNWGRISNVLPEFRGEGGV    240
                *********************************** :****************:***.**

Stx2a_A         RVGRISFNNISAILGTVAVILNCHHQGARSVRAVNEESQPECQITGDRPVIKINNTLWES    300
Stx2dact_A      RVGRISFNNISAILGTVAVILNCHHQGARSVRAVNEEIQPECQITGDRPVIRINNTLWES    300
                ************************************* *************:********

Stx2a_A         NTAAAFLNRKSQFLYTTGK*    319
Stx2dact_A      NTAAAFLNRRAHSLNTSGE*    319
                *********::: * *:*:*
```

```
Stx2a_B         MKKMFMAVLFALASVNAMAADCAKGKIEFSKYNEDDTFTVKVDGKEYWTSRWNLQPLLQS    60
Stx2dact_B      MKKIFVAALFAFVSVNAMAADCAKGKIEFSKYNENDTFTVKVAGKEYWTNRWNLQPLLQS    60
                ***:*:*.***:.*********************:******* ******.**********

Stx2a_B         AQLTGMTVTIKSSTCESGSGFAEVQFNND*  89
Stx2dact_B      AQLTGMTVTIKSNTCASGSGFAEVQFN*--  87
                ************.** ***********
```

**Figure S3: Stx2 alignment.** Alignment of the A and B subunits of Stx2a and Stx2dact generated using Clustal Omega.

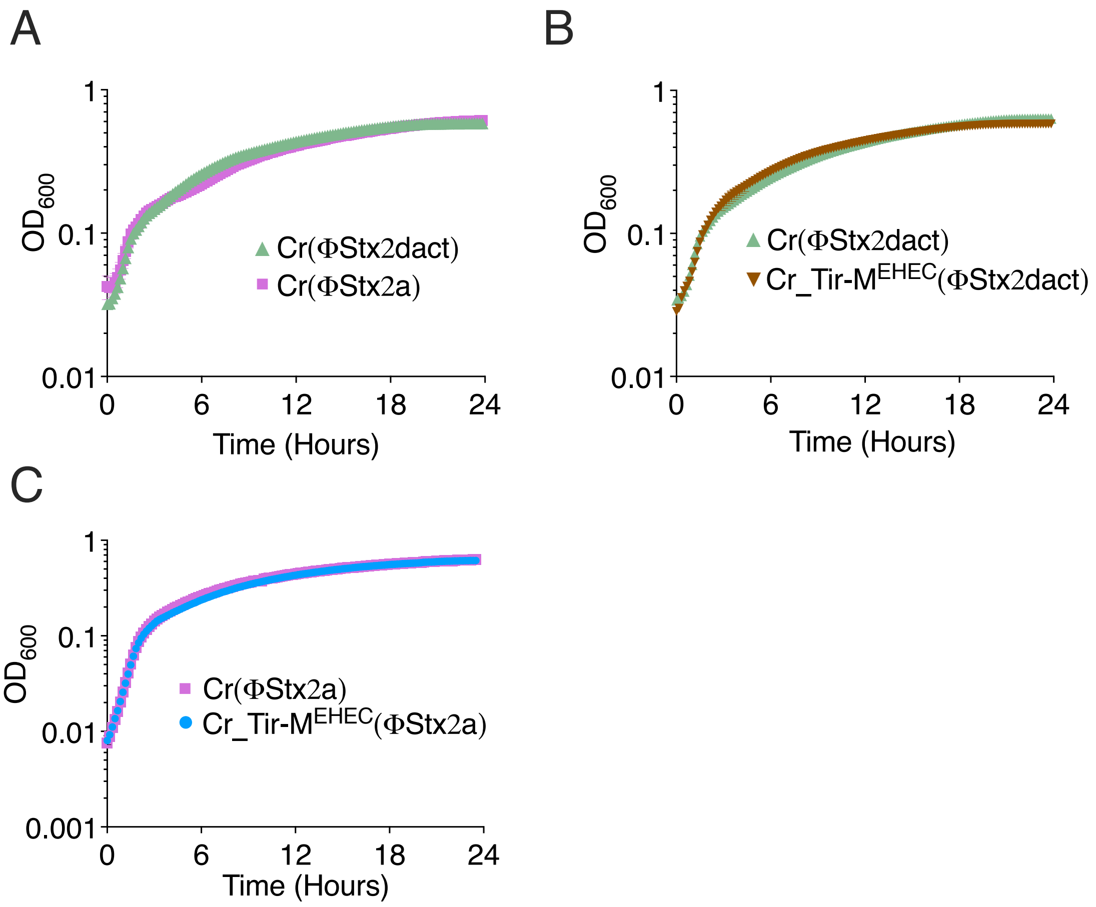


**Figure S4: Growth curves of Cr(FStx2) based strains.** (A-C) Growth curves of Cr(ΦStx2dact) vs. Cr(ΦStx2a) (A), Cr(ΦStx2dact) and Cr_Tir-M$^{EHEC}$(ΦStx2dact) (B) and Cr(ΦStx2a) and Cr_Tir-M$^{EHEC}$(ΦStx2a) (C). The data shown is representative of the mean $\underline{+}$ SEM of four technical repeats.

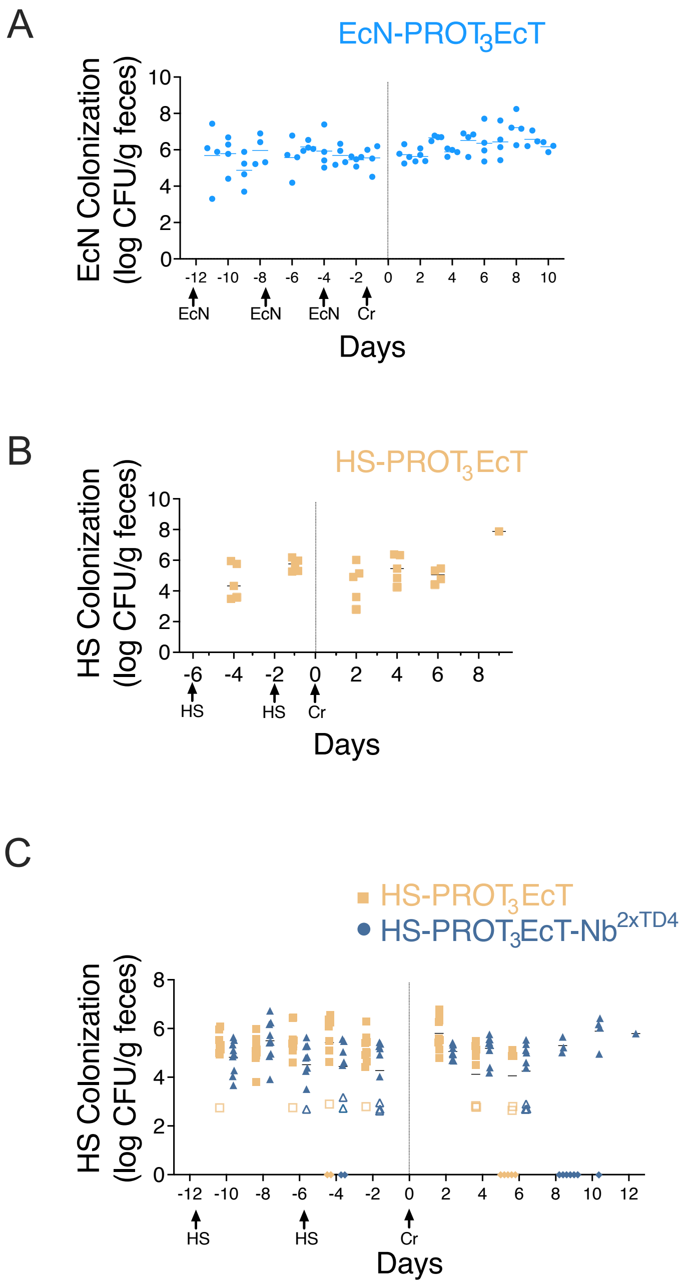


**Figure S5: EcN- and HS-PROT3EcT colonization of mice.** (A, B, C) Titers of fecally shed bacteria from mice inoculated with EcN- (A) or HS-PROT3EcT (B, C). Arrows denote when EcN-PROT3EcT, HS-PROT3EcT, or Cr(Stx2) was administered. Five mice are included in each cohort. Each point shown represents an individual mouse, and each line represents the geometric mean.

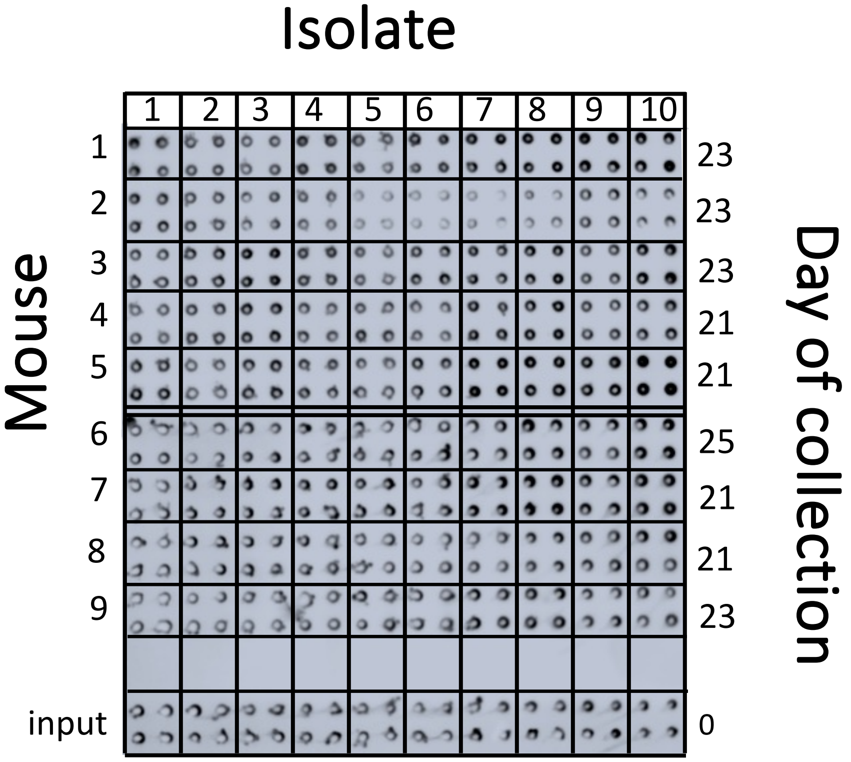


**Figure S6: HS-PROT$_3$EcT-TD4 maintains activity for at least 21 days.** Plate secretion assay of quad-spotted single colonies isolated from each of the 9 mice pre-treated with HS-PROT$_3$EcT-Nb$^{2xTD4}$. The 10 colonies studied were each obtained from the last day shed feces from the designated mouse.

**Table S1.** Plasmids

| Plasmid | Description | Reference |
|---|---|---|
| pTKRED | λ-Red recombinase/SceI endonuclease expression plasmid, $SPEC^R$ | (1) |
| pDONR221 | Entry plasmid containing a Gateway cassette, pUC ori, $KAN^R$ | Thermo Fisher Scientific |
| pCP20 | Thermal induction of FLP recombinase, temperature sensitive, $AMP^R$ | (2) |
| pKD4 | template plasmid for FRT-flanked KAN cassette, R6K ori, KANR | (2) |
| pDonor-Stx2a | Donor plasmid carrying Stx2a locus, pDONR221 derivative, $KAN^R$ | This study |
| pDonor-Cr-Tir-$M^{EHEC}$ | Donor plasmid carrying Cr-Tir-$M^{EHEC}$ , pDONR221 derivative, $KAN^R$ | This study |
| pDSW206 | IPTG-inducible expression vector, ColE1 ori, $AMP^R$ | (3) |
| pDSW206-ccdB-FLAG | pDSW206 derivative containing a Gateway cassette + rrnB terminator, $AMP^R$, $CM^R$ | (4) |
| pCPG-alr | DSW206 derivative containing EcN *alr* under endogenous promoter + T7 terminator, ColE1 ori, $AMP^R$ | (5) |
| pTKIP-PJ23119-virB | *virB* under J23119 synthetic promoter flanked by LP and SceI, $AMP^R$, $HYGRO^R$ | (5) |
| pDSW206-$SS^{OspC2}$-$Nb^{1xTNF}$ | Expression construct, p*tac* (IPTG) *ospC2ss-*$Nb^{1xTNF}$*-3xFLAG*, ColE1 origin, $AMP^R$ | (5) |
| pDSW206-$SS^{OspC2}$-$Nb^{2xTNF}$ | Expression construct, p*tac* (IPTG) *ospC2ss-*$Nb^{2xTNF}$*3xFLAG*, ColE1 origin, $AMP^R$ | (5) |
| pENTR221-$SS^{OspC2}$-$Nb^{1xTD4}$ | Entry plasmid, *ospC2SS-1xNb*$^{TD4}$, pUC ori, $KAN^R$ | This study |
| pENTR221-$SS^{OspC2}$-$Nb^{2xTD4}$ | Entry plasmid, *ospC2SS-2xNb*$^{TD4}$, pUC ori, $KAN^R$ | This study |
| pDSW206-$SS^{OspC2}$-$Nb^{1xTD4}$ | Expression construct, ptac (IPTG) *ospC2ss-*$Nb^{1xTD4}$*-3xFLAG*, ColE1 origin, $AMP^R$ | This study |
| pDSW206-$SS^{OspC2}$-$Nb^{2xTD4}$ | Expression construct, ptac (IPTG) ospC2ss-$Nb^{2xTD4}$-3xFLAG, ColE1 origin, $AMP^R$ | This study |
| pCPG-alr-$SS^{OspC2}$-$Nb^{2xTD4}$ | Expression construct, PJ23108 *ospC2ss-*$Nb^{2xTD4}$-3xFLAG, ColE1 origin, $AMP^R$ | This study |
| pCPG-alr-$SS^{OspC2}$-$Nb^{2xTD4}$ | Expression construct, PJ23108 *ospC2ss-*$Nb^{2xTNF}$-3xFLAG, ColE1 origin, $AMP^R$ | (5) |

**Table S2.** Strains

| Strain | Description | Reference |
|---|---|---|
| **Citro strains** | | |
| Cr(ΦStx2dact) | DBS770 | (6) |
| Cr(ΦStx2::Tet$^{R}$) | Cr(ΦStx2dact) with SceI-TetR-SceI swapped with Stx2dact | This study |
| Cr(ΦStx2a) | DB3770 with Stx2a | This study |
| Cr_Tir-M$^{TetR}$(ΦStx2dact) | Cr(ΦStx2dact) with SceI-TetR-SceI swapped with Tir-M | This study |
| Cr_Tir-M$^{TetR}$(ΦStx2a) | Cr(ΦStx2a) with SceI-TetR-SceI swapped with Tir-M | This study |
| Cr_Tir-M$^{EPEC}$(ΦStx2a) | Cr(ΦStx2da) with EPEC Tir-M domain | This study |
| Cr_Tir-M$^{EHEC}$(ΦStx2a) | Cr(ΦStx2da) with EPEC Tir-M domain | This study |
| **PROT3EcT strains** | | |
| T3EcT | DH10b with *mxi-spa* operons | (5) |
| EcN-PROT$_3$EcT | PROT$_3$EcT-4 | (5) |
| PROT$_3$EcT-1- LP$^{yie/trk}$ | PROT$_3$EcT-1 with LP at yieN/trkB locus | (5) |
| PROT$_3$EcT-2 | HS with Mxi-Spa operons at atp1/gidB locus | (5) |
| PROT$_3$EcT-2-LP$^{yie/trk}$ | PROT$_3$ECT-2 with LP at yieN/trkB locus | This study |
| PROT$_3$EcT-2-virB | PROT$_3$ECT-2 with J23119-virB at yieN/trkB locus | This study |
| PROT$_3$EcT-2-virB ΔdadX::kanR | PROT3EcT-2-virB with ΔdadX::kan | This study |
| PROT$_3$EcT-2-virB ΔdadX | PROT3EcT-2-virB with ΔdadX | This study |
| HS-PROT$_3$EcT | PROT3EcT-2-virB ΔdadxΔalr::kan | This study |
| HS-PROT$_3$EcT-TD4 | HS-PROT$_3$EcT with pCPG-alr-SS$^{OspC2}$-Nb$^{2xTD4}$ | This study |

**Table S3 Primers**

| Primer | Sequence |
|---|---|
| **oligos for generating Cr-based strains** | |
| P1 | cctgcgcccggccctttagc |
| P2 | gaaaggtggtgtcaacgtaaatATTACCCTGTTATCCCTAAAACTCCCGGGAATAGGATA |
| P3 | AACAGGAGACAAGTGCTTAGTAGGGATAACAGGGTAATtatcagaagcattgctggtttc |
| P4 | ggccacgcagttgcgcagag |
| P5 | TAGGGATAACAGGGTAATatttacgttgacaccacctttc |
| P6 | ATTACCCTGTTATCCCTACTAAGCACTTGTCTCCTGTTTA |
| P7 | gcgctggggcgtcgtgtaac |
| P8 | ggcgctggggcgtcgtgt |
| P9 | aggccacgcagttgcgca |
| P10 | accgggggacgtggtggtag |
| P11 | aaggtggtgtcaacgtaaatATTACCCTGTTATCCCTAcgcctgaacaatacctgtagct |
| P12 | TAAACAGGAGACAAGTGCTTAGTAGGGATAACAGGGTAATtcatcaggtattggttacgg |
| P13 | GGAGCGAGAAGCGATGGATTATTcctgg |
| P14 | TTAATTCCGCCTGCGCCGCC |
| P15 | GACGGCGCGACAAGGGGG |
| P16 | CGCAGCGCTCCCTGCGCT |
| **oligos for generating HS-PROT$_3$EcT-based strains** | |
| P17 | CTTTTAGCGCCTGAATAGAA |
| P18 | TGGTACCAATATCGCCGTA |
| P19 | CAATAAGCTTAAAAGGCCATCCGTCAGGA |
| P20 | ACGTCATTTAAGTAGCAGTTAAGGCTGTTTTGGCGGATGAG |
| P21 | ggccatttacatggcgcacacagctaaggaaacgagatgacccgtgtgtaggctggagctgcttc |
| P22 | gtggattaatcgttctgtaatatttgattgtctgtgccggcatatgaatatcctccttag |
| P23 | aattaaagcaaacacttatcaaggaacacaaatgcaagcggtgtaggctggagctgcttc |
| P24 | ccggcacagacaatcaaatattacagaacgattaatccaccatatgaatatcctccttag |
| P25 | CCGCTGAAAGGCTACTCGCT |
| P26 | CGGTTGCGATGCTTTGCTG |
| P27 | CCGTTCTCTGGAACAACGTGC |
| P28 | GCCAAGGGTTTGAATGGTTGC |

**Table S4 Synthetic DNA fragment sequences**

| **attB1-Stx hybrid-attB2** |
| --- |
| AAGGTCGTCAAAATGGGTGTCGTAATTTATCAGCGCGACAACTTTGTACAAAAAAGTTGGC<br>TAGGGATAACAGGGTAATttcaaatccagcaagggccaccatatcacataccgccattagctcatcgggatagagcgcc<br>agccttcgaagctggctgcgcggggttcgagtcctcgatggcggtccattatcggtattcagcgttgttagctcagccggacagagca<br>attgccttctaagcaatcggtcactggttcgaatccagtacaacgcgccatacttattttttctGGCTCGCTTTTGCGGGCCT<br>TTTTTATATCTGCGCCGGGTCTGGTGCTGATTACTTCAGCCAAAAGGAACACCTGTATatgaa<br>gtgtatattatttaaatgggtactgtgcctgttactgggtttttcttcggtatcctattcccgggagtttACGATAGACTTTTCGACC<br>CAACAAAGTTATGTCTCTTCGTTAAATAGTATACGGACAGAGATATCGACCCCTCTTGAACAT<br>ATATCTCAGGGGACCACATCGGTGTCTGTTATTAACCACACCCCACCGGGCAGTTATTTTG<br>CTGTGGATATACGAGGGCTTGATGTCTATCAGGCGCGTTTTGACCATCTTCGTCTGATTATT<br>GAGCAAAATAATTTATATGTGGCCGGGTTCGTTAATACGGCAACAAATACTTTCTACCGTTTT<br>TCAGATTTTACACATATATCAGTGCCCGGTGTGACAACGGTTTCCATGACAACGGACAGCA<br>GTTATACCACTCTGCAACGTGTCGCAGCGCTGGAACGTTCCGGAATGCAAATCAGTCGTC<br>ACTCACTGGTTTCATCATATCTGGCGTTAATGGAGTTCAGTGGTAATACAATGACCAGAGAT<br>GCATCCAGAGCAGTTCTGCGTTTTGTCACTGTCACAGCAGAAGCCTTACGCTTCAGGCAG<br>ATACAGAGAGAATTTCGTCAGGCACTGTCTGAAACTGCTCCTGTGTATACGATGACGCCGG<br>GAGACGTGGACCTCACTCTGAACTGGGGGCGAATCAGCAATGTGCTTCCGGAGTATCGG<br>GGAGAGGATGGTGTCAGAGTGGGGAGAATATCCTTTAATAATATATCAGCGATACTGGGGA<br>CTGTGGCCGTTATACTGAATTGCCATCATCAGGGGGCGCGTTCTGTTCGCGCCGTGAATG<br>AAGAGAGTCAACCAGAATGTCAGATAACTGGCGACAGGCCTGTTATAAAAATAAACAATACA<br>TTATGGGAAAGTAATACAGCTGCAGCGTTTCTGAACAGAAAGTCACAGTTTTTATATACAAC<br>GGGTAAATAAaggagttaagcATGAAGAAGATGTTTATGGCGGTTTTATTTGCATTAGCTTCTGTT<br>AATGCAATGGCGGCGGATTGTGCTAAAGGTAAAATTGAGTTTTCCAAGTATAATGAGGATGA<br>CACATTTACAGTGAAGGTTGACGGGAAAGAATACTGGACCAGTCGCTGGAATCTGCAACC<br>GTTACTGCAAAGTGCTCAGTTGACAGGAATGACTGTCACAATCAAATCCAGTACCTGTGAA<br>TCAGGCTCCGGATTTGCTGAAGTGCAGTTTAATAATGACTGAtatcagaagcattgctggtttcgtggtgtg<br>cagcaatgtagttacagtgtaatcaatgtcacaattcagtcagttgaaggttgtctgcccgactgagaatttgttaaaaaaaaaatcctg<br>catggtgaatccccctgagcggcggggcatatcagcgtcacaggtgtttctgttttacctctatcctttctgtgcgggttcaggtgctgata<br>ctgaactcaccgggaggcacccggcaccatgcatgaacggtacatagcgcatacatcagcccctctccggaggggctttcttgtgg<br>gcaaaaaaaaTAGGGATAACAGGGTAATCCAACTTTCTTGTACAAAGTTGTGCCGACAATATT<br>CTGTCTAGCTTGGCGCTAGC |
| **Stx hybrid_TET PCR fragment** |
| cctgcgcccggccctttagctcagtggtgagagcgagcgactcataatcgccaggtcgctggTTCAAATCCAGCAAGGG<br>CCACCATATCACATACCGCCATTAGCTCATCGGGATAGAGCGCCAGCCTTCGAAGCTGGCT<br>GCGCGGGGTTCGAGTCCTCGATGGCGGTCCATTATCGGTATTCAGCGTTGTTAGCTCAGC<br>CGGACAGAGCAATTGCCTTCTAAGCAATCGGTCACTGGTTCGAATCCAGTACAACGCGCC<br>ATACTTATTTTTTCTGGCTCGCTTTTGCGGGCCTTTTTTATATCTGCGCCGGGTCTGGTGCT<br>GATTACTTCAGCCAAAAGGAACACCTGTATATGAAGTGTATATTATTTAAATGGGTACTGTGC<br>CTGTTACTGGGTTTTTCTTCGGTATCCTATTCCCGGGAGTTTTAGGGATAACAGGGTAATattt<br>acgttgacaccacctttcgcgtatggcatgatagcgcccggaagagagtcaattcagggtggtgaatATGAATAGTTCGAC<br>AAAGATCGCATTGGTAATTACGTTACTCGATGCCATGGGGATTGGCCTTATCATGCCAGTCT<br>TGCCAACGTTATTACGTGAATTTATTGCTTCGGAAGATATCGCTAACCACTTTGGCGTATTGC<br>TTGCACTTTATGCGTTAATGCAGGTTATCTTTGCTCCTTGGCTTGGAAAAATGTCTGACCGA<br>TTTGGTCGGCGCCCAGTGCTGTTGTTGTCATTAATAGGCGCATCGCTGGATTACTTATTGCT<br>GGCTTTTTCAAGTGCGCTTTGGATGCTGTATTTAGGCCGTTTGCTTTCAGGGATCACAGGA<br>GCTACTGGGGCTGTCGCGGCATCGGTCATTGCCGATACCACCTCAGCTTCTCAACGCGTG<br>AAGTGGTTCGGTTGGTTAGGGGCAAGTTTTGGGCTTGGTTTAATAGCGGGGCCTATTATTG<br>GTGGTTTTGCAGGAGAGATTTCACCGCATAGTCCCTTTTTTATCGCTGCGTTGCTAAATATT<br>GTCACTTTCCTTGTGGTTATGTTTTGGTTCCGTGAAACCAAAAATACACGTGATAATACAGAT<br>ACCGAAGTAGGGGTTGAGACGCAATCAAATTCGGTGTACATCACTTTATTTAAAACGATGCC<br>CATTTTGTTGATTATTTATTTTTCAGCGCAATTGATAGGCCAAATTCCCGCAACGGTGTGGG<br>TGCTATTTACCGAAAATCGTTTTGGATGGAATAGCATGATGGTTGGCTTTTCATTAGCGGGT<br>CTTGGTCTTTTACACTCAGTATTCCAAGCCTTTGTGGCAGGAAGAATAGCCACTAAATGGG<br>GCGAAAAAACGGCAGTACTGCTCGGATTTATTGCAGATAGTAGTGCATTTGCCTTTTTAGCG<br>TTTATATCTGAAGGTTGGTTAGTTTTCCCTGTTTTAATTTTATTGGCTGGTGGTGGGATCGCT |

| |
|---|
| TTACCTGCATTACAGGGAGTGATGTCTATCCAAACAAAGAGTCATCAGCAAGGTGCTTTACA<br>GGGATTATTGGTGAGCCTTACCAATGCAACCGGTGTTATTGGCCCATTACTGTTTGCTGTTA<br>TTTATAATCATTCACTACCAATTTGGGATGGCTGGATTTGGATTATTGGTTTAGCGTTTTACTG<br>TATTATTATCCTGCTATCAATGACCTTCATGTTGACCCCTCAAGCTCAGGGGAGTAAACAGG<br>AGACAAGTGCTTAGTAGGGATAACAGGGTAATtatcagaagcattgctggtttcgtggtgtgcagcaatgtagtta<br>cagtgtaatcaatgtcacaattcagtcagttgaaggttgtctgcccgactgagaatttgttaaaaaaaaaatcctgcatggtgaatccc<br>cctgagcggcggggcatatcagcgtcacaggtgtttctgttttacctctatcctttctgtgcgggttcaggtgctgatactgaactcaccgg<br>gaggcacccggcaccatgcatgaacggtacatagcgcatacatcagcccctctccggaggggctttcttgtgggcaaaaaaaaa<br>aagcccgcgccgggagacgcgggcggcaaggaataaacaacaaaacgtgaagtaaatttcagctggcgaataatacccgaca<br>gtaatcactctgcgcaactgcgtggcc |
| **attB1-EHEC_Cr_Tir-M-attB2** |
| CAGGGGGACAACTTTGTACAAAAAAGTTGGCTAGGGATAACAGGGTAATgatggtacccatgctgct<br>attggggtaaaagatggtgtagaggttagcgttacattaaattctagtgaattgcaaagcttgcaatctcttgacactgaaggtaacggt<br>cgatttgttttcaccgggggacgtggtggtagtgggcatgccatggtcactgttgcatcagatatctcgcaagctcgtgagaaaataat<br>agctaagttagatccagacaaccatggaggaagacaacctaaggacattgatacacgttctgttggtgtcggaagtgcttcaggaat<br>gggagatggagttgttagcgaaactcatacttcaacaacaacttcaagtgttcgttcagatcctaaattctgggtttctgttggcgcaatt<br>gctgctggtttagcggggctggcagctacaggtattgttcaggcgcttgcattgacgccggagccggatagcccaaccacgaccga<br>ccctgatgcagctgcaagtgcaactgaaactgcgacaagagatcagttaacgaaagaagcgttccagaacccagataatcaaaa<br>agttaatatcgatgagctcggaaatgcgattccgtcaggggtattgaaagatgatgttgttgcgaatatagaagagcaggctaaagc<br>agcaggcgaagaggccaaacagcaagccattgaaaataatgctcaggcgcaaaaaaaatatgatgaacaacaagctaaacg<br>ccaggaggagctgaaagtttcatcaggtattggttacggcctcagtagtgcattgattgttggtggggggattggtgctggtgttacggc<br>tatgctccatagacgaaatccgccgacagaacagacaatagctactacacattcggttattcagcagcagaccggggaaatact<br>cgagcacaaggcggcgctgacaccactggagtggaaaacgcttctctgactagacgtgattcgcaggctagtgttgcatcgaccca<br>atggtcagatacctctggcgatgtggtgaatccgtatgctgaaggttggatgtccaggaataatccatcgcttctcgctccagaagagc<br>ctatttatgatgaagtcgctccggatcctaactatagcgttattcagcatttttcagggaataatcctgtgactgggcggttagtaggaTA<br>GGGATAACAGGGTAATCCAACTTTCTTGTACAAAGTTGTCCCCGAC |
| **attB1-SS$^{OspC2}$-Nb$^{1xTD4}$-attB2** |
| CGAGGGGACAACTTTGTACAAAAAAGTTGGCGAAGGAGATAGAACCATGAAAATACCTGAA<br>GCAGTAAATCATATTAATGTTCAGAACAATATTGATCTTGTTGATGGAAAAATAAATCCGAACA<br>AAGATACAAAAGCATTACAAAAAAACATATCATGCGTAACAAACTCATCCTCTTCTGGCATAA<br>GTGAAAAAGGTGGAGGTGGCTCCGGCGGCGGAGGTTCCCAGGTCCAGTTGGTAGATGCC<br>GGTGGTGGCAGCGTGCAAGCTGGGGGGTCGCTTACACTTTCCTGTGTGGCCTCCGGCGC<br>AGCGTATAGTACTAATCTTCTGGGATGGTTCCGCCAGGCCCCTGGGAAAGAGCGCGAAGG<br>GGTTGCTTCTATCTACCGTGGCAATAGCGCAACAAATTACGCTGATTCAGTCAAGGGCCGC<br>TTTACCATCAGCCAAGACAAAACGAAGTACACCATTTACCTGCAGATGAACTCTCTTAAACC<br>CGAGGACAGTGCTATGTACTACTGCGCACACGGTACAGCCCCATATTGGCACACCCCGAT<br>CCCAACTTTATCAGAGGATAAGTACTTCTATTGGGGACAGGGCACACAAGTGACCGTGTCC<br>TCTCCAACTTTCTTGTACAAAGTTGTCCCCGACC |
| **attB1-SS$^{OspC2}$-Nb$^{2xTD4}$-attB2** |
| GGGACAAGTTTGTACAAAAAAGTTGGCGGAATTCCTGACAGCTAGCTCAGTCCTAGGTATA<br>ATGCTAGCTGATAAATGCTTCAATAATATTGAAAAAGGAGGAGTATGAAAATACCTGAAGCA<br>GTAAATCATATTAATGTTCAGAACAATATTGATCTTGTTGATGGAAAAATAAATCCGAACAAAG<br>ATACAAAAGCATTACAAAAAAACATATCATGCGTAACAAACTCATCCTCTTCTGGCATAAGTG<br>AAAAAGGCGGTGGTGGTTCGGGTGGTGGCGGCTCTCAGGTCCAACTCGTAGATGCAGGC<br>GGTGGTTCCGTTCAGGCCGGCGGGAGCTTAACCTTGAGTTGTGTGGCCTCTGGCGCAGC<br>GTACAGTACTAACTTACTCGGTTGGTTTCGACAGGCACCAGGAAAGGAACGTGAGGGTGT<br>GGCATCGATCTACCGTGGTAATTCGGCAACGAATTATGCGGATTCAGTGAAAGGTAGATTTA<br>CAATTAGCCAAGACAAGACTAAATACACAATTTATCTGCAAATGAACTCTCTGAAGCCGGAA<br>GACTCAGCCATGTACTATTGTGCTCACGGGACTGCACCGTACTGGCACACACCAATACCTA<br>CTTTGAGTGAGGACAAATACTTTTATTGGGGTCAGGGAACCCAAGTGACGGTTAGTTCGG<br>GTGGCGGCGGCTCAGGTGGAGGAGGCAGCCAAGTACAGCTCGTTGATGCAGGAGGAGG<br>CTCCGTGCAGGCAGGTGGAAGTCTGACTTTGTCCTGCGTTGCGTCAGGCGCGGCTTACT<br>CCACTAATTTGTTAGGTTGGTTCCGGCAGGCCCCCGGGAAGGAACGGGAAGGCGTGGCT<br>AGCATTTATCGCGGAAATTCTGCCACTAATTACGCAGATTCGGTGAAGGGGAGATTTACCAT<br>CAGTCAGGATAAGACTAAATATACTATATATTTGCAGATGAATAGCTTAAAGCCAGAAGACTC<br>AGCTATGTATTATTGCGCGCACGGGACCGCACCGTACTGGCATACCCCTATACCGACTCTG |

AGCGAAGACAAATATTTCTACTGGGGACAAGGGACACAAGTTACCGTTTCATCTAGACCAA
CTTTCTTGTACAAAGTTGTCCCCGACCGGG

**SI References**


1. T. E. Kuhlman, E. C. Cox, Site-specific chromosomal integration of large synthetic constructs. *Nucleic Acids Res* **38**, e92 (2010).
2. K. A. Datsenko, B. L. Wanner, One-step inactivation of chromosomal genes in *Escherichia coli* K-12 using PCR products. *Proc. Natl. Acad. Sci. U.S.A.* **97**, 6640–6645 (2000).
3. D. S. Weiss, J. C. Chen, J.-M. Ghigo, D. Boyd, J. Beckwith, Localization of FtsI (PBP3) to the Septal Ring Requires Its Membrane Anchor, the Z Ring, FtsA, FtsQ, and FtsL. *J Bacteriol* **181**, 508–520 (1999).
4. A. M. Schmitz, M. F. Morrison, A. O. Agunwamba, M. L. Nibert, C. F. Lesser, Protein interaction platforms: visualization of interacting proteins in yeast. *Nat Methods* **6**, 500–502 (2009).
5. J. P. Lynch, *et al.*, Engineered *Escherichia coli* for the in situ secretion of therapeutic nanobodies in the gut. *Cell Host & Microbe* **31**, 634-649.e8 (2023).
6. E. M. Mallick, *et al.*, A novel murine infection model for Shiga toxin–producing Escherichia coli. *J. Clin. Invest.* **122**, 4012–4024 (2012).
